# A Data-Driven Transcriptional Taxonomy of Adipogenic Chemicals to Identify White and Brite Adipogens

**DOI:** 10.1101/519629

**Authors:** Stephanie Kim, Eric Reed, Stefano Monti, Jennifer Schlezinger

## Abstract

**Background:** Chemicals in disparate structural classes activate specific subsets of PPARγ’s transcriptional programs to generate adipocytes with distinct phenotypes.

**Objectives:** Our objectives were to 1) establish a novel classification method to predict PPARγ ligands and modifying chemicals, and 2) create a taxonomy to group chemicals based on their effects on PPARγ’s transcriptome and downstream metabolic functions. We tested the hypothesis that environmental adipogens highly ranked by the taxonomy, but segregated from therapeutic PPARγ ligands, would induce white but not brite adipogenesis.

**Methods:** 3T3-L1 cells were differentiated in the presence of 76 chemicals (negative controls, nuclear receptor ligands known to influence adipocyte biology, potential environmental PPARγ ligands). Differentiation was assessed by measuring lipid accumulation. mRNA expression was determined by RNA-Seq and validated by RT-qPCR. A novel classification model was developed using an amended random forest procedure. A subset of environmental contaminants identified as strong PPARγ agonists were analyzed by their effects on lipid handling, mitochondrial biogenesis and cellular respiration in 3T3-L1 cells and human preadipocytes.

**Results:** We used lipid accumulation and RNA sequencing data to develop a classification system that 1) identified PPARγ agonists, and 2) sorted chemicals into likely white or brite adipogens. Expression of *Cidec* was the most efficacious indicator of strong PPARγ activation. Two known environmental PPARγ ligands, tetrabromobisphenol A and triphenyl phosphate, which sorted distinctly from therapeutic ligands, induced white adipocyte genes but failed to induce *Pgc1a* and *Ucp1*, and induced fatty acid uptake but not mitochondrial biogenesis in 3T3-L1 cells. Moreover, two chemicals identified as highly ranked PPARγ agonists, tonalide and quinoxyfen, induced white adipogenesis without the concomitant health-promoting characteristics of brite adipocytes in mouse and human preadipocytes.

**Discussion:** A novel classification procedure accurately identified environmental chemicals as PPARγ ligands distinct from known PPARγ-activating therapeutics. The computational and experimental framework has general applicability to the classification of as-yet uncharacterized chemicals.

## Introduction

Since 1980, the prevalence of obesity has been increasing in the United States (Flegal et al. 2016). Further, in 2015, it was estimated that a total of 108 million children and 604 million adults were obese worldwide (GBD 2017). This poses a major public health threat since overweight and obesity increase the risk of metabolic syndrome, which, in turn, sets the stage for metabolic diseases, such as type 2 diabetes, cardiovascular disease, nonalcoholic fatty liver disease and stroke (Park et al. 2003). The Endocrine Society’s latest scientific statement on the obesity pathogenesis states that obesity is a disorder of the energy homeostasis system, rather than just a passive accumulation of adipose, and that environmental factors, including chemicals, confer obesity risk (Schwartz et al. 2017). The rapid increases in obesity and metabolic diseases correlate with substantial increases in environmental chemical production and exposures over the last few decades, and experimental evidence in animal models demonstrates the ability of a broad spectrum of various environmental metabolism-disrupting chemicals to induce adiposity and metabolic disruption (Heindel et al. 2017).

Adipocytes are essential for maintaining metabolic homeostasis as they are the storage depot of free fatty acids and release hormones that can modulate body fat mass (Rosen and Spiegelman 2006). Adipogenesis is a highly regulated process that involves a network of transcription factors acting at different time points during differentiation (Farmer 2006).

Peroxisome proliferator activated receptor γ (PPARγ) is a ligand activated, nuclear receptor, which is required for adipocyte formation and function (Tontonoz et al. 1994), as well as metabolic homeostasis. In both PPARγ haploinsufficient (Gumbilai et al. 2016) and knockout (He et al. 2003; Jiang et al. 2014; O’Donnell et al. 2016; Zhang et al. 2004) rodent models, there was a lack of adipocyte formation and metabolic disruption.

PPARγ regulates energy homeostasis by activating expression of genes involved in both storage of excess energy as lipids in white adipocytes and energy utilization by triggering mitochondrial biogenesis, fatty acid oxidation and thermogenesis in brite (brown-in-white) and brown adipocytes. The white adipogenic, brite/brown adipogenic and insulin sensitizing activities of PPARγ are regulated distinctly through ligand-specific post-translational modifications (Banks et al. 2015; Choi et al. 2010; Choi et al. 2011; Qiang et al. 2012) and co-regulator recruitment (Burgermeister et al. 2006; Feige et al. 2007; Ohno et al. 2012; Villanueva et al. 2013). While adult humans have long been thought to not have brown adipose tissue, humans with minimal brite adipocyte populations are at higher risk for obesity and type 2 diabetes (Claussnitzer et al. 2015; Sidossis and Kajimura 2015; Timmons and Pedersen 2009).

Growing evidence supports the hypothesis that environmental PPARγ ligands induce phenotypically distinct adipocytes. Tributyltin (TBT) induces the formation of an adipocyte with lower adiponectin expression and altered glucose homeostasis (Regnier et al. 2015).

Furthermore, TBT failed to induce expression of genes associated with browning of adipocytes (e.g. *Ppara*, *Pgc1a*, *Cidea*, *Elovl3*, *Ucp1*) in differentiating 3T3-L1 adipocytes (Kim et al. 2018; Shoucri et al. 2018). As a result, TBT-induced adipocytes failed to up-regulate mitochondrial biogenesis and had low levels of cellular respiration (Kim et al. 2018; Shoucri et al. 2018). The structurally similar environmental PPARγ ligand, triphenyl phosphate, also fails to induce brite adipogenesis, and this correlates with an inability to prevent PPARγ from being phosphorylated at Serine 273 (S273) (Kim et al. 2020).

The EPA developed the Toxicity Forecaster (ToxCast™) program to use high-throughput screening assays to prioritize chemicals and inform regulatory decisions regarding thousands of environmental chemicals (Kavlock et al. 2012). Several ToxCast™ assays can measure the ability of chemicals to bind to or activate PPARγ, and these assays have been used to generate a toxicological priority index (ToxPi) that were expected to predict the adipogenic potential of chemicals in cell culture models (Auerbach et al. 2016). Yet, it was shown that the results of ToxCast™ PPARγ assays do not always correlate well with activity measured in a laboratory setting and that the ToxPi designed for adipogenesis was prone to predicting false positives (Janesick et al. 2016). Furthermore, the ToxCast/ToxPi approach cannot distinguish between white and brite adipogens (Pereira-Fernandes et al. 2014).

In this study, we investigate differences in cellular response between adipogenic and non-adipogenic compounds, as well as the heterogeneity of response across adipogenic compounds. Our ultimate goal is to create a method for identification of novel adipogenic compounds using the taxonomic organization of known and predicted adipogenic compounds based on their divergent transcriptional response. To this end, we generated phenotypic and transcriptomic data from adipocytes differentiated in the presence of 76 different chemicals. We combined the cost-effective generation of agonistic transcriptomic data by 3’Digital Gene Expression – a highly multiplexed RNA-Seq technology – with a new classification method to predict PPARγ-activating and modifying chemicals. Further, we investigated metabolic-related outcome pathways as effects of the chemical exposures. We created a data-driven taxonomy to specifically classify chemicals into distinct categories based on their various interactions with and effects on adipogenesis, presumably through their interaction with PPARγ. Based on the taxonomy-based predictions, we tested the phenotype (white *vs.* brite adipocyte functions) of environmental adipogens predicted to fail to induce brite adipogenesis in 3T3-L1 cells and primary human adipocytes.

## Methods

### Chemicals

DMSO was purchased from American Bioanalytical (Natick, MA). CAS numbers, sources and catalog numbers of experimental chemicals are provided in **Table S1**. Human insulin, dexamethasone, 3-isobutyl-1-methylxanthine (IBMX), and all other chemicals were from Sigma-Aldrich (St. Louis, MO) unless noted.

**Table 1.** Summary of experimental conditions

|  | 3T3-L1 | HUMAN<br>PREADIPOCYTES |
| --- | --- | --- |
| Exposure Period (days) | 10 | 14 |
| Number of times dosed | 4 | 6 |
| Positive Control | Rosiglitazone | Rosiglitazone |
| Negative Control (Vehicle) | DMSO | DMSO |

### Cell Culture

3T3 L1 (RRID:CVCL_0123, Lot # 63343749, ATCC, Manassas, VA) cells were originally derived from a Swiss mouse embryonic fibroblast line (Green and Kehinde, 1975). Cells were maintained in Growth Medium (high-glucose DMEM (Corning, 10-013-CV) with 10% calf serum (Sigma), 100 U/ml penicillin, 100 μg/ml streptomycin, 0.25 μg/ml amphotericin B). All experiments were conducted with cells between passages 3 and 9. Experimental conditions are outlined in **Table 1** and **Figure S1A**. For experiments, cells were plated in Growth Medium and incubated for 4 days, at which time the cultures are confluent for 2 days. “Naïve” pre-adipocytes were cultured in every experiment and grown in Growth Medium for the duration of an experiment. On day 0, differentiation was induced by replacing the medium with Differentiation Medium (DMEM,10% fetal bovine serum (FBS, Sigma-Aldrich), 100 U/ml penicillin, 100 μg/ml streptomycin, 250 nM dexamethasone (Figures 1-3), 1 nM dexamethasone (Figures 5-10) or no dexamethasone (Figure S3), 167 nM human insulin, 0.5 mM IBMX). Also on day 0, single experimental wells were treated with vehicle (DMSO, 0.2% final concentration), rosiglitazone (positive control, 200 nM) or test chemicals. On days 3 and 5 of differentiation, medium was replaced with Maintenance Medium (DMEM, 10% FBS, 167 nM human insulin, 100 U/ml penicillin, 100 μg/ml streptomycin), and the cultures were re-dosed. On Day 7 of differentiation, medium was replaced with Adipocyte Medium (DMEM, 10% FBS, 100 U/ml penicillin, 100 μg/ml streptomycin), and the cultures were re-dosed. On day 10, cytotoxicity was assessed by microscopic inspection, with cultures containing more than 10% rounded cells excluded from consideration. For most chemicals only a single concentration was tested; however, for those chemicals inducing toxicity the concentration was reduced until no toxicity was observed (See **Table S1** for information on maximum tested and maximum non-toxic concentrations tested). Wells with healthy cells were harvested for analysis of gene expression, lipid accumulation, fatty acid uptake, mitochondrial biogenesis, mitochondrial membrane potential, and cellular respiration.

**Figure 1.**
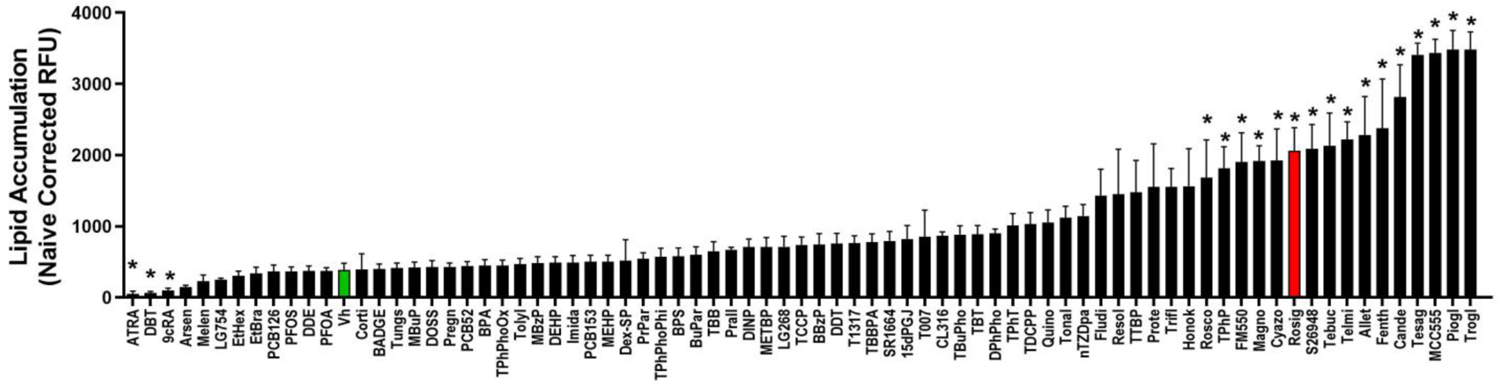
Lipid accumulation in differentiated and treated 3T3-L1 pre-adipocytes. Confluent 3T3 L1 cells were differentiated using a standard hormone cocktail for 10 days. During differentiation, cells were treated with vehicle (Vh, 0.2% DMSO, final concentration), rosiglitazone (100 nM) or test chemicals (**Table S1**). On days 3, 5, and 7 of differentiation, the medium was replaced and the cultures re-dosed. Following 10 days of differentiation and dosing, cells were analyzed for lipid accumulation by Nile Red staining. Nile Red fluorescence was normalized by subtracting the fluorescence measured in naïve pre-adipocyte cultures within each experiment and reported as “Naïve Corrected RFU.” Numerical data are provided in **Excel File 3**. Data are presented as mean ± SE (n=4). Statistically different from Vh-treated (highlighted in green) (*p<0.05, ANOVA, Dunnett’s).

OP9 cells (RRID:CVCL_4398, Lot # 63544739, ATCC) are a bone marrow stromal cell line derived from newborn calvaria of the (C57BL/6xC3H)F2-op/op mouse (Nakano *et al*., 1994). Cells were maintained in αMEM (Gibco, 12-561-056) with 20% FBS, 26 mM sodium bicarbonate, 100 U/ml penicillin, 100 μg/ml streptomycin, 0.25 μg/ml amphotericin B. Cells were plated in 24 well plates at 50,000 cells were well in 500 µl medium and incubated for 4 days. Induction and maintenance of adipogenesis and treatment were as described for 3T3 L1 cells, except that the dexamethasone concentration was 125 nM.

Primary human subcutaneous pre-adipocytes from five individual female patients were obtained from the Boston Nutrition Obesity Research Center (Boston, MA). The patients were 32-51 years of age and had BMIs ranging from 26.0-30.7. One patient was pre-diabetic, and four were non-diabetic. Adipocytes were differentiated as previously described (Lee and Fried 2014). Experimental conditions are outlined in **Table 1** and **Figure S1B**. The pre-adipocytes were maintained in Growth Medium (αMEM with 10% FBS, 100 U/ml penicillin, 100 μg/ml streptomycin, 0.25 μg/ml amphotericin B). For experiments, human pre-adipocytes were plated in Growth Medium and grown to confluence (3-5) days. “Naïve” pre-adipocytes were cultured in every experiment and grown in Growth Medium for the duration of an experiment. On day 0, differentiation was induced by replacing the Growth Medium with Differentiation Medium (DMEM/F12, 25 mM NaHCO_3_, 100 U/ml penicillin, 100 μg/ml streptomycin, 33 μM d-Biotin, 17 μM pantothenate, 100 nM dexamethasone, 100 nM human insulin, 0.5 mM IBMX, 2 nM T_3_, 10 μg/ml transferrin). Also on day 0, single experimental wells also were treated with vehicle (DMSO, 0.1% final concentration), rosiglitazone (positive control, 4 μM) or test chemicals. On day 3 of differentiation, medium was replaced with fresh Differentiation Medium, and the cultures were re-dosed. On days 5, 7, 10, and 12 of differentiation, the medium was replaced with Maintenance Medium (DMEM/F12, 25 mM NaHCO_3_, 100 U/ml penicillin, 100 μg/ml streptomycin, 3% FBS, 33 μM d-Biotin, 17 μM pantothenate, 10 nM dexamethasone, 10 nM insulin), and the cultures were re-dosed. Following 14 days of differentiation and dosing, cells were harvested for analysis of gene expression, lipid accumulation, fatty acid uptake, mitochondrial biogenesis, and cellular respiration.

### Lipid Accumulation

3T3-L1 cells or human preadipocytes were plated in 24 well plates at 50,000-100,000 cells per well in 0.5 ml maintenance medium at initiation of the experiment. Dosing is outlined in **Table 1**. Medium was removed from the differentiated cells, and they were rinsed with PBS. The cells were then incubated with Nile Red (1 µg/ml in PBS) for 15 min in the dark. Fluorescence (λex= 485 nm, λem= 530 nm) was measured using a Synergy2 plate reader (BioTek Inc., Winooski, VT). The fluorescence in all experimental wells was normalized by subtracting the fluorescence measured in naïve pre-adipocyte cultures within each experiment and reported as “Naïve Corrected RFU.”

### Transcriptome Profiling

3T3-L1 cells were plated in 24 well plates at 50,000 cells per well in 0.5 ml maintenance medium at initiation of the experiment. Dosing is outlined in **Table 1**. Total RNA was extracted and genomic DNA was removed using the Direct-zol MagBead RNA Kit and following manufacturer’s protocol (Zymo Research, Orange, CA). RNA concentrations and contamination were determined spectrophotometrically using a Nanodrop (ND-1000, ThermoFisher). A final concentration of 5 ng RNA/ul was used for each sample. For each chemical, 3-5 biological replicates were profiled and carefully randomized across six 96-well plates, including 26 DMSO vehicle controls, and 16 naïve pre-adipocyte cultures. Sequencing and gene expression quantification was carried out by the MIT Technology Lab at Broad Institute (Cambridge, MA). RNA libraries were prepared using a highly multiplexed 3’ Digital Gene Expression (3’ DGE) protocol developed by (Xiong et al. 2017) and sequenced on an Illumina NextSeq 500, generating between 2.13E8 and 3.87E8 reads and a mean of 3.02E8 reads per lane across 96 samples. All reads containing bases with Phred quality scores < Q10 were removed. The remaining reads were aligned to mouse reference genome, GRCm38, and counted in 21,511 possible transcripts annotations. Only instances of uniquely aligned reads were quantified (i.e., reads that aligned to only one transcript). Furthermore, multiple reads with the same unique molecular identifier (UMI), aligning to the same gene were quantified as a single count.

### Reverse Transcription (RT)-qPCR

3T3-L1 cells or human preadipocytes were plated in 24-well plates at 100,000 cells per well in 0.5 ml maintenance medium at initiation of the experiment. Dosing is outlined in **Table 1**. Total RNA was extracted and genomic DNA was removed using the 96-well Direct-zol MagBead RNA Kit (Zymo Research). RNA concentrations and contamination were determined spectrophotometrically using a Nanodrop. cDNA was synthesized from total RNA using the iScript™ Reverse Transcription System (BioRad, Hercules, CA). All qPCR reactions were performed using the PowerUp™ SYBR Green Master Mix (Thermo Fisher Scientific, Waltham, MA). The qPCR reactions were performed using a 7500 Fast Real-Time PCR System (Applied Biosystems, Carlsbad, CA): UDG activation (50°C for 2 min), polymerase activation (95°C for 2 min), 40 cycles of denaturation (95°C for 15 sec) and annealing (various temperatures for 15 sec), extension (72°C for 60 sec). The primer sequences and annealing temperatures are provided in **Table S2**. All primers were obtained from Integrated DNA Technologies (Coralville, IA).

**Table 2.**
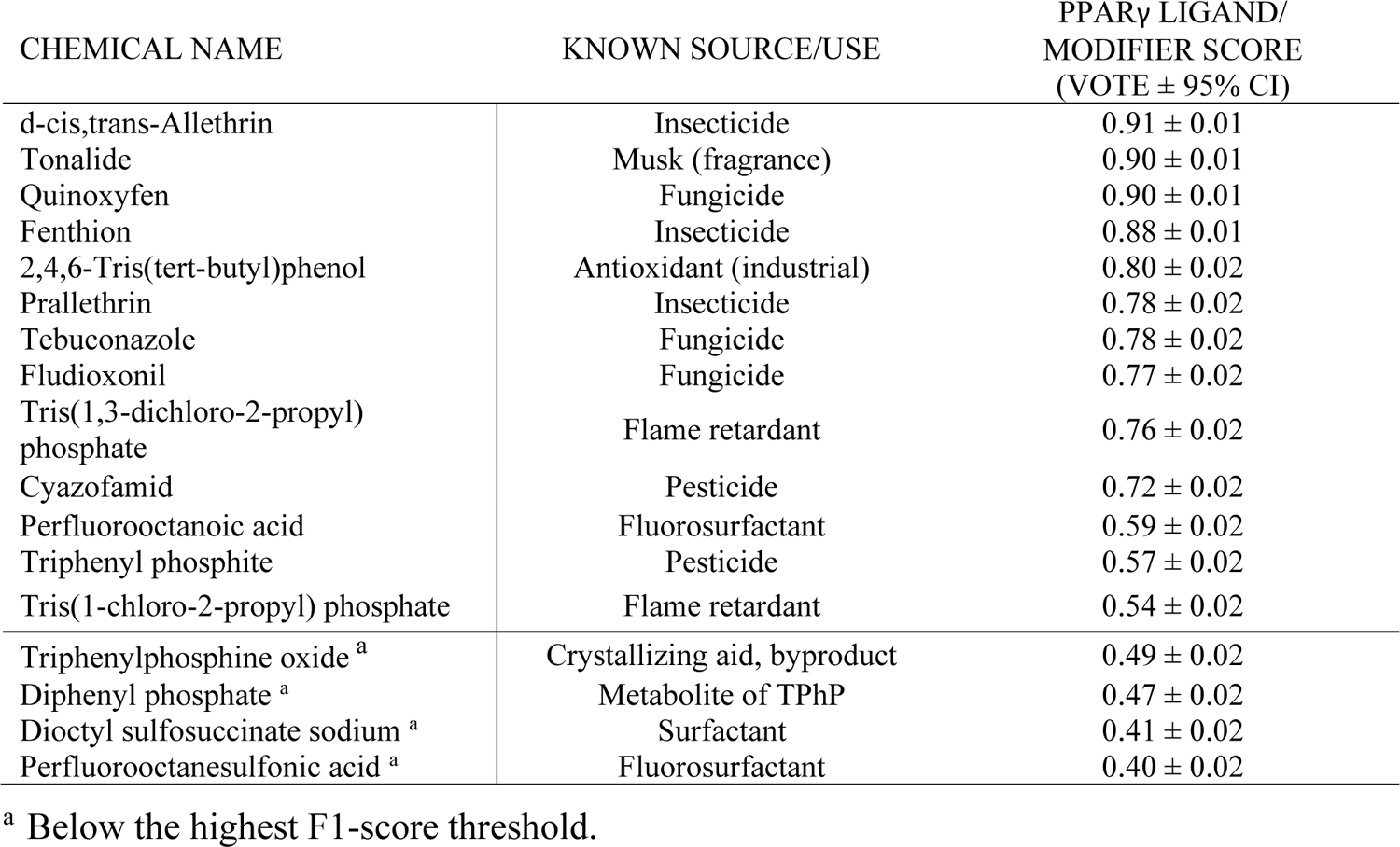
Amended random forest classification results for 17 compounds that are potential PPARγ Ligands/Modifiers.

Relative gene expression was determined using the Pfaffl method to account for differential primer efficiencies (Pfaffl 2001), using the geometric mean of the Cq values for beta-2-microglobulin (*B2m*) and 18s ribosomal RNA (*Rn18s*) for mouse gene normalization and of ribosomal protein L27 (*RPL27*) and *B2M* for human gene normalization. The Cq value from naïve, pre-adipocyte cultures within each experiment was used as the reference point for the experiment. Data are reported as “Relative Expression.”

### Cell Number Analysis

3T3-L1 cells or human preadipocytes were plated in 96-well, black-sided plates at 12,500 cells per well in 0.2 ml maintenance medium at initiation of the experiment. Dosing is outlined in **Table 1**. Cellular protein was measured by Janus Green staining and measuring absorbance (595 nM) using a Synergy2 plate reader. The absorbance in experimental wells was normalized by dividing by the absorbance measured in naïve pre-adipocyte cultures within the experiment and reported as “Relative Cell Density.”

### Fatty Acid Uptake

3T3-L1 cells or human preadipocytes were plated in 96-well, black-sided plates at 12,500 cells per well in 0.2 ml maintenance medium at initiation of the experiment. Dosing is outlined in **Table 1**. Fatty acid uptake was measured by treating differentiated cells with 100 μL of Fatty Acid Dye Loading Solution (Sigma-Aldrich, MAK156). Fluorescence intensity (λex= 485nm, λem= 530nm) measured at time zero and after a 10-minute incubation using a Synergy2 plate reader. The fluorescence at time zero was subtracted from the fluorescence at the end of the incubation and as “RFU.”

### Mitochondrial Biogenesis

3T3-L1 cells or human preadipocytes were plated in 24-well plates at 100,000 cells per well in 0.5 ml maintenance medium at initiation of the experiment. Dosing is outlined in **Table 1**. Mitochondrial biogenesis was measured in differentiated cells using the MitoBiogenesis In-Cell Elisa Colorimetric Kit, following the manufacturer’s protocol (Abcam). The expression of the mitochondrial protein SDH was measured and normalized to total protein content via Janus Green staining. Absorbance (OD 405nm for SDH, and OD 595nm for Janus) was measured using a Synergy2 plate reader. The absorbance ratios of SDH/Janus are reported as “Relative Mitochondrial Protein Expression.”

### Oxygen Consumption

3T3-L1 cells were plated in 24-well plates at 100,000 cells per well in 0.5 ml maintenance medium at initiation of the experiment. Dosing is outlined in **Table 1**. After 10 days of differentiation, cells were gently trypsinized, diluted in 400 ul Adipocyte Medium, and 80 ul per well was transferred to duplicate wells of a 96-well Agilent Seahorse plate. After 24 hrs incubation, the medium was changed to Seahorse XF Assay Medium without glucose (1mM sodium pyruvate, 1mM GlutaMax, pH 7.4) and cultures incubated at 37°C in a non-CO_2_ incubator for 1 hr. To measure mitochondrial respiration, the Agilent Seahorse XF96 Cell Mito Stress Test Analyzer (available at BUMC Analytical Instrumentation Core) was used, following the manufacturer’s standard protocol. The compounds and their concentrations used to determine oxygen consumption rate (OCR) included 1) 0.5 μM oligomycin, 1.0 μM carbonyl cyanide-p-trifluoromethoxyphenylhydrazone (FCCP) and 2 μM rotenone/2 μM antimycin. After the Seahorse analysis, cells were fixed in 4% paraformaldehyde and stained with Janus Green.

Respiration rates were normalized by dividing by the Janus Green absorbance and reported as “Relative OCR.” Basal respiration was determined by subtracting non-mitochondrial respiration from the last rate measurement before the injection of oligomycin. Maximum respiration was determined by subtracting non-mitochondrial respiration from the maximum rate measurement after the injection of FCCP. Spare capacity was determined by subtracting the basal respiration from the maximum respiration.

### Adiponectin Secretion

3T3-L1 cells were plated in 24-well plates at 100,000 cells per well in 0.5 ml maintenance medium at initiation of the experiment. Dosing is outlined in **Table 1**. After 9 days of differentiation, the medium was replaced with washout medium (DMEM+0.1% BSA), and cultures were incubated for 24 hrs. The washout medium was replaced, the cells redosed and the incubation continued for 24 hrs. Samples of medium were collected, diluted 1:200 and analyzed for adiponectin by ELISA following the manufacturer’s instructions (Mouse Adiponectin/Acrp30 Quantikine ELISA Kit, R&D Systems, MRP300). Absorbance (450 nM) was measured using a Synergy2 plate reader and concentrations calculated relative to a standard curve.

### Statistical Analyses

All statistical analyses were performed in R (v 3.4.3) and Prism 7 (GraphPad Software, Inc., La Jolla, CA). All R code used for processing and analysis of transcriptome profiles is publicly available on GitHub (https://github.com/montilab/Adipogen2020) and were carried out using several R packages. Normalization and differential gene expression analysis was performed using *limma* (v 3.34.9) (Ritchie et al. 2015). Batch correction was performed using *ComBat* (v 3.26.0) (Leek et al. 2012). Gene set projection was performed using *GSVA* (v 1.30.0) (Hanzelmann et al. 2013). Statistical testing of partial correlation estimates was performed using *ppcor* (v 1.1) (Kim 2015).

For 3T3-L1 experiments the biological replicates correspond to independently plated experiments. For human primary preadipocyte experiments the biological replicates correspond to distinct individuals’ preadipocytes (5 individuals in all). For each experiment, naïve, undifferentiated 3T3-L1 cells were included for within-experiment normalization. The Nile Red data and the qPCR data were not normally distributed; therefore, the data were log transformed before statistical analyses. One-factor ANOVAs (Dunnett’s) were performed to analyze the qPCR and phenotypic data and determine differences from vehicle-treated cells. Data are presented as means ± standard error (SE).

### Transcriptome Data Processing

The number of counted reads per sample transcriptome profile varied widely with a range of 7.90E1 to 2.27E6 (Mean = 2.25E5, SD = 2.94E5). To remove technical noise introduced by low overall expression quantification of individual samples, we performed an iterative clustering-based approach to determine sets of samples which segregate as a result of low total read counts. Each iteration included four steps: removal of low count genes, normalization, plate-level batch correction, and hierarchical clustering. Low count genes, defined as having mean counts <1 across all samples, were removed to reduce statistical noise introduced by inaccurate quantification of consistently lowly expressed transcripts. Normalization was performed using Trimmed Mean of M-values (TMM), the default method employed by *limma* (v 3.34.9) (Ritchie et al. 2015). Batch correction was performed by *ComBat* (v 3.26.0) (Leek et al. 2012).

Hierarchical clustering was performed on the 3000 genes with the largest median absolute deviation (MAD) score, using Euclidean distance and 1-Pearson correlation as the distance metric for samples and genes, respectively, and Ward’s agglomerative method (Ward 1963). Clusters of samples clearly representative of low expression quantification were removed. This process was repeated until no such low expression outlier sample cluster was present (four iterations). For the remaining samples, low count genes were removed, and samples were normalized and batch corrected by the same procedure.

Following sample- and gene-level quality control filtering, the final processed data set included expression levels of 9,616 genes for each of 234 samples. These 234 samples included 2-4 remaining replicates of each compound, 25 DMSO vehicle controls, and 15 naïve pre-adipocyte cultures. Sequencing data from 3’DGE have been deposited to GEO (Accession: GSE124564).

### PPARγ Activating/Modifier Classification

A classification model was inferred from the *training set* consisting of 37 known PPARγ-modifying compounds and 23 known non-PPARγ modifying compounds, including vehicle, to predict the label of the *test set* of 17 potential PPARγ-modifying compounds (**Table S1**). Known compounds were selected based on a literature search for experimental evidence of modification of PPARγ activity, including PPARγ binding assays, coactivator recruitment or computational modeling (definitive ligands), PPARγ-driven reporter assays (at least 25% of the rosiglitazone-induced maximum), expression of PPARγ target genes and/or differentiation of 3T3 L1 or multipotent stromal cells into adipocytes in the absence of a known PPARγ ligand (**Table S1**). Known non-PPARγ modifying compounds were selected based on a literature search for chemicals that influence adipogenesis but that are ligands for other nuclear receptors (e.g., aryl hydrocarbon receptor, glucocorticoid receptor) or selected based on their structural dissimilarity from known PPARγ ligands and lack of evidence of PPARγ activation (**Table S1**). The model inference was based on an amended random forest procedure developed to better account for the presence of biological replicates in the data. Specifically, for each classification tree, samples and genes were bagged using sampling techniques consistent with (Breiman 2001). In particular, samples were bootstrapped (i.e., sampled with replacement), and genes were subsampled by the square root of the number of represented genes. To account for chemical-level variability and to prevent replicates of the same chemical exposure from being separated, we implemented an extra step, which we denote as “bag-merging”, to summarize values from replicate samples of the same chemical exposure after bootstrap sampling: within each “bag” of samples, replicates of the same chemical exposure were merged to their mean expression. For prediction of test data, replicates of the same chemical exposure were merged to their mean expression and run through the trained random forest, such that each tree generates a vote of either 0 or 1, PPARγ-modifying negative or positive, respectively. The mean of these votes across all trees is a value between 0 and 1, which can be interpreted as the pseudo-probability of the chemical exposure being PPARγ-modifying.

Prior to generating the final predictive model, the expected performance of this classification approach for predicting PPARγ-modification status with this data set was assessed using 10-fold cross validation on the training set of known PPARγ-modification status. For each fold, samples were stratified at the chemical exposure level, such that each fold included 6 distinct compounds, and all replicates of the same compound were only included in either the training or the test folds. Next, prior to training each random forest model, gene filtering based on between *vs.* within exposure variance using ANOVA was performed. Genes with an F-statistics associated with an FDR corrected p-value > 0.05 were filtered out. Thresholds for determining class membership based on voting was determined by running the training folds through the random forest and selecting the threshold producing the highest F1-score, i.e., the harmonic mean of precision and sensitivity. Performance was assessed in terms of area under the curve (AUC), as well as precision, sensitivity, specificity, F1-score, and balanced accuracy, i.e., the mean of specificity and sensitivity. All random forests were generated using 2000 decision trees.

The performance of this procedure was compared to three alternative random forest strategies. In the first, which we denote as “pre-merging”, the mean gene expression across replicates was computed, and a classic random forest was applied to the classification of each merged chemical profile. In the second, which we denote as “classic”, replicate samples were treated as independent perturbations and classified based on a classic random forest. Finally, for the third, which we denote as “pooled”, the mean of votes across replicates from the “classic” strategy were used to estimate class membership per compound. To compare the classifiers’ performance, we repeated the 10-fold CV procedure 10-times to generate a distribution of performance statistics for each strategy.

The final classification model used to predict the unlabeled chemicals was built using the full training set of 59 labelled chemicals, including vehicle, and 1,199 genes after performing the same ANOVA gene filtering approach used for cross validation as described above. The full list of genes used in this model, including chemical-specific log fold-changes, F-statistics, nominal P-values and FDR corrected P-values are provided in **Excel File 1**. The relative importance of each gene in each random forest model was measured using the Gini importance, which evaluates the mean decrease of label impurity when a particular gene is used to separate chemicals in a given tree (Breiman 2001). Maximum purity is achieved (Gini=0) when all chemicals within a branch have the same label.

### Adipogen Clustering

Known and potential PPARγ activators/modifiers were clustered based on their test statistics from univariate analysis comparing each chemical exposure or naïve pre-adipocyte culture to vehicle using *limma* (v 3.34.9) (Ritchie et al. 2015). In order to assess taxonomic differences between different exposure outcomes, a recursive unsupervised procedure, which we denote as “*K2Taxonomer*” (Reed et al. 2020), was developed, whereby the set of PPARγ modifying compounds underwent recursive partitioning into subgroups. At each iteration of the procedure, the top 10% of genes was selected based on estimation of non-random changes in gene expression across the current set or subset of compounds via the sum of squared test statistics. Chemicals were then separated into two clusters based on their Euclidean distance, using Ward’s agglomerative method (Ward 1963) and the *cutree* R function. The procedure was recursively applied to each of the two identified subgroups of chemicals and repeated until the two-cluster split would result in a single chemical in the terminal subgroup. To obtain and measure the most stable clusters, each iteration was bootstrapped 200 times by resampling gene-level statistics with replacement. The most common clusters were used, and the proportion of total bootstrapping iterations that included these identical clustering assignments was reported.

In order to derive gene-signatures of each split, differential analysis was performed to compare chemicals from the two clusters at that split. In these models, biological replicate status was accounted for using the duplicate correlation procedure using *limma* (v 3.34.9) (Ritchie et al. 2015). From these models, differential signatures were defined whereby genes were assigned to one of four genesets based on two criteria: their differential expression between the two chemical clusters at the split, and – within each chemical cluster – the differential expression between chemicals and vehicle. In particular, for a particular gene, the difference between mean expression between the two chemical clusters must have |log2(Fold-Change)|> 1 (i.e., Fold-Change>2) and an FDR Q-value < 0.1. Each gene was then assigned to one-of-four genesets: 1) genes with higher expression in the “left” chemical group vs. the “right” chemical group, and with higher expression in the left chemical group vs. vehicle; 2) genes with higher expression in the left chemical group vs. the right chemical group, and lower expression in the left chemical group vs. vehicle; 3) genes with higher expression in the right chemical group vs. the left chemical group, and with higher expression in the right chemical group vs. vehicle; 2) genes with higher expression in the right chemical group vs. the left chemical group, and with lower expression in the right chemical group vs. vehicle. Since the results of direct differential analysis between the two chemical clusters are not indicative of overall up- or down-regulation, these designations were determined based on the aggregate of the comparisons of each chemical or naïve pre-adipocyte culture to vehicle. Specifically, a gene was assigned to a cluster based on maximum absolute value of the mean of the chemical or naïve pre-adipocyte culture versus vehicle derived test statistics used for clustering. Direction of regulation was then determined based on the sign of the mean of these test statistics. Functional enrichment, comparing these gene sets to independently annotated gene sets was carried out via Fisher’s exact test. These gene sets include those of the full set of Gene Ontology (GO) Biological Processes gene set compendia downloaded from MSigDB, c5.bp.v6.2.symbols.gmt (Subramanian et al. 2005; Liberzon et al. 2011), as well two gene sets derived from publicly available microarray expression data from an experiment using mouse embryonic fibroblasts to compare wild-type samples with mutant samples that do not undergo phosphorylation of PPARγ at Ser273, GEO accession number GSE22033 (Choi et al. 2010). These additional gene sets were comprised of genes, measured to be significantly higher or lower in expression (FDR Q-Value < 0.05) in mutant samples, based on differential analysis of RMA normalized expression with *limma* (v 3.34.9). The full set of differential gene expression analysis and functional enrichment analysis results, including the chemicals belonging to each subgroup, are provided in **Excel File 2**.

### Human Transcriptome Analysis

To assess the human relevance of the gene signatures derived from the adipogen clustering in chemical-exposed 3T3-L1 cells, we analyzed projections of these signatures onto the transcriptional profiles from 770 subjects who were part of the Metabolic Syndrome in Men (METSIM) study. Although comprised of only male subjects, the METSIM data set was chosen for this analysis because it is the largest publicly available human subject data set, GEO accession number GSE70353 (Civelek et al. 2017), which includes gene expression profiles from subcutaneous adipose tissue, as well as a comprehensive set of cardio-metabolic measurements. Affymetrix Human Genome U219 Array Microarray CEL files were annotated to unique Entrez gene IDs, using a custom CDF file from BrainArray (HGU219_Hs_ENSG_22.0.0). Each of the four possible gene sets derived from adipogen clustering were then projected on each of the 770 human transcriptome profiles using gene set variation analysis (GSVA), using *GSVA* (v 1.30.0) (Hanzelmann et al. 2013), resulting in an enrichment score for each gene set and sample.

For each projected gene set, we tested for relationships between single-sample enrichment scores and clinical measurements. Of notice, many of the clinical measurements are correlated with each other, such that confounding is likely to generate many spurious results. To overcome this problem, for each gene set projection, we tested the significance of the partial correlation between single-sample enrichment scores and each of the clinical variables while controlling for the remaining ones, including age, using *ppcor* (v 1.1) (Kim 2015). Given that the relationships between single-sample enrichment scores and any one clinical variable are not assumed to be linear, these partial correlations were calculated from Spearman correlation estimates. P-values were adjusted across all combinations of gene set and clinical measurement. Measurements with at least one comparison with a gene set projection yielding an FDR Q-value < 0.1 are reported. To reduce expected redundancies in the measurements, of the 23 initial quantitative measurements included in these data, we ran this analysis on a subset of 12 measurements (**Table S3**). Of note, fat free mass % was chosen in this subset of 12 measurements over body mass index because it has been shown to be better associated with risk and presence of metabolic syndrome (Liu et al. 2013).

## Results

### Classification of novel taxonomic subgroups of adipogens

Potential adipogens (chemicals that change the differentiation and/or function of adipocytes) were identified by review of the literature for reports of PPARγ agonism or modulation of adipocyte differentiation or by the ToxPi designed to identify chemicals in the ToxCast dataset that have potential to be PPARγ ligands/modifiers (**Table S1**). Our classification groups were labeled as “Yes”, “No”, or “Potential”, based on the chemical’s potential ability to act as a ligand of or modify PPARγ (i.e., to alter its post translational modifications) as noted in the “PPARγ Ligand or Modifier” column in **Table S1**.

The classic mouse pre-adipocyte model, 3T3-L1 cells, was differentiated and treated with vehicle (DMSO) or with each of the 76 test chemicals 4 times over the ten-day course of differentiation (concentrations are reported in **Table S1**). In order to maximize the number of chemicals that could be characterized, each chemical was tested at a single, maximal, non-toxic dose. We also limited the maximum concentrations to 20 μM because concentrations above this would not be reached in humans and because most (although not all) chemicals are not toxic at or below 20 μM. Lipid accumulation was determined after 10 days. Lipid accumulation spanned from significantly lower to significantly higher than vehicle-treated cells (**Figure 1**). **Excel File 3** contains numerical data from all cell culture experiments. Lipid accumulation was highly correlated with expression of adipocyte specific genes (e.g., *Cidec* and *Fabp4* (Danesch et al. 1992), **Figure S2**); therefore, we considered lipid accumulation a biomarker of adipocyte differentiation in this system. Of the 27 chemicals that significantly induced lipid accumulation, 18 were known PPARγ ligands/modifiers and 9 were potential PPARγ ligands/modifiers (**Figure 1**, **Table S1**). Nine of the 17 potential adipogens generated significantly higher differentiation than Vh-treated cells, including Alleth, Cyazo, Fenth, Fludiox, Quinox, TDCPP, Tebu, Tona, and TTBP.

We turned to two other models to investigate why these differences from other studies could be occurring. First, we differentiated the 3T3 L1 cells in the presence of the same suite of chemicals, but in the absence of dexamethasone in the differentiation cocktail. The adipogenic effects of steroids (e.g., melengesterol, cotisosterone, dexamethasone) were significant when dexamethasone was not included in the cocktail (**Figure S3a**). Although, adipogenesis in the presence and absence of 250 nM dexamethasone was moderately correlated overall (**Figure S3b**). Second, we differentiated OP9 cells, a late preadipocyte model (Wolins et al. 2006), in the presence of the same suite of chemicals. OP9 cells were more sensitive to differentiation stimulated by RXR ligands (**Figure S4a**). However, adipocyte differentiation stimulated in OP9 cells was still moderately correlated with differentiation stimulated in 3T3 L1 cells differentiated with 250 nM dexamethasone (**Figure S4b**).

Following the analysis of lipid accumulation, RNA was isolated from the cells, and the transcriptome was characterized by RNA-Seq. The transcriptome data were then used to develop a classification model to identify adipogens likely to act as PPARγ activators/modifiers. More specifically, the *training set* of 59 chemicals with “Yes”/“No” labels was used to build the classifier, which was then applied to the prediction of the *test set* of 17 “Potential” ligands. When predicting PPARγ activation/modifier status (“Yes” *vs.* “No”), the mean area-under-the-curve (AUC), precision, sensitivity, specificity, F1-score, and balanced accuracy from repeated 10-fold cross validation (over the training set) of the random forest with bag merging procedure was 0.87, 0.85, 0.79, 0.76, 0.82, and 0.77, respectively (**Figure 2A**). We observed the most drastic improvement of measured balanced accuracy, precision, and specificity by the bag merging procedure compared to other assessed strategies (**Figure S5**). The full set of cross-validation results for each performance metrics and all 10 repetitions can be found in **Excel File 1**. The first two metrics in particular (AUC and precision) reflect expectation of relatively few false positive results compared to the other strategies. In the final model fitted on the entire training set, the voting threshold that produced the highest F1-score was 0.53. When we applied the classifier to the test set of the 17 chemicals of unknown interaction with PPARγ, 13 had random forest votes greater than this value (**Table 3**). Of these 13 compounds, four had votes > 0.88. These chemicals included quinoxyfen, tonalide, allethrin, and fenthion. These compounds were predicted as PPARγ activators/modifiers with high confidence and were selected for further functional analyses. Of the 1,215 genes that passed ANOVA filtering and were included in the random forest models, ribosomal protein L13 (*Rpl13*) and cell death Inducing DFFA Like Effector C (*Cidec*) were measured to have the highest Gini importance (**Figure 2B**); *Rpl13* was primarily expressed at a lower level and *Cidec* was primarily expressed at a higher level in cells treated with known PPARγ activators/modifiers (**Figure S6**). Gene specific ANOVA filtering results and importance estimates, as well as out-of-bag voting estimates for each compound in the training set can be found in **Excel File 1**.

**Figure 2.**
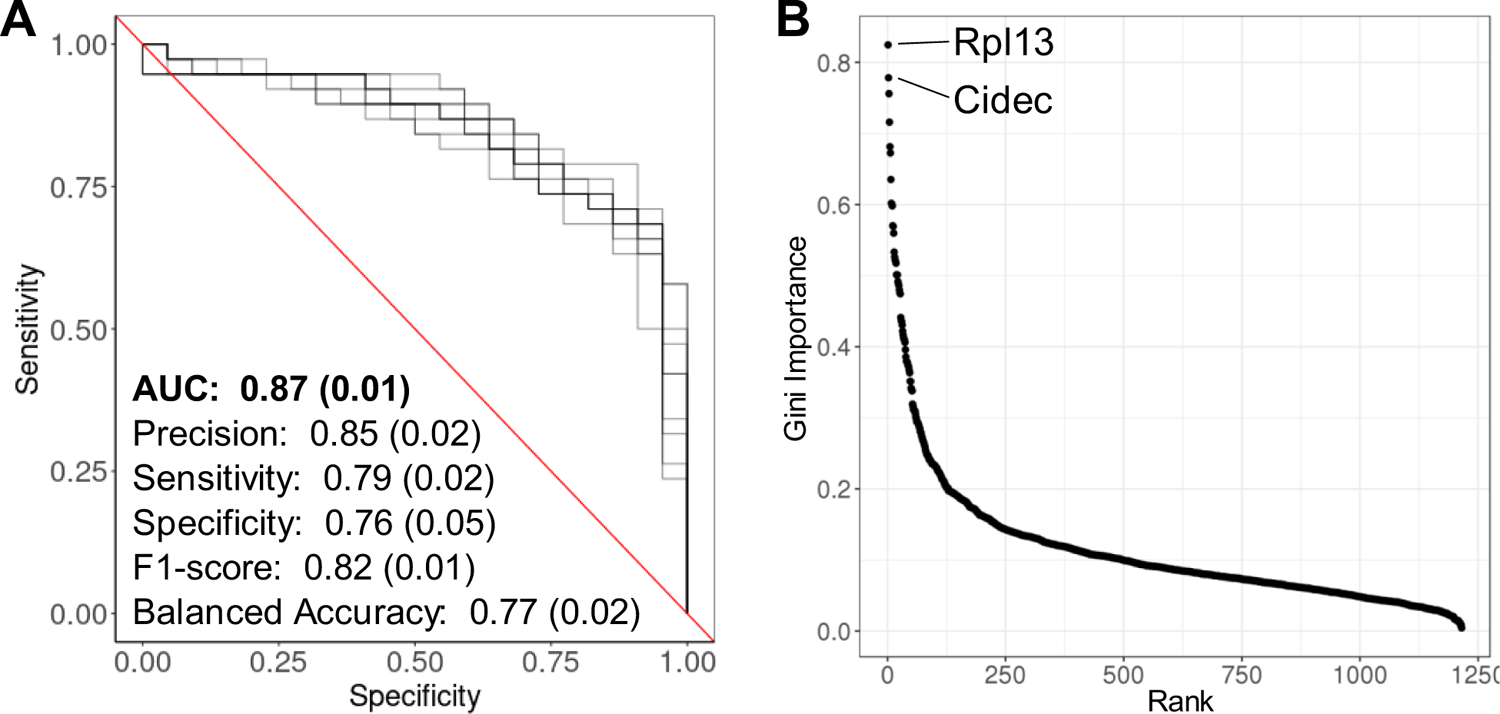
Amended random forest classification performance and gene importance of final classification model. **(A)** Performance of random forest classification procedure based 10-fold cross validation. The plot shows the receiver operating characteristic curves associated with the AUC for 10 repetitions of performance estimation. The values represent the mean estimate across repetitions for each evaluation metric. The standard deviations these distributions are shown in parenthesis. The full set of performance estimates for each repetition, performance metric, and classification procedure considered are shown in **Excel File 1**. **(B)** Gini importance versus ranking of genes used in the final random forest model. The names of the top 2 genes are highlighted. Compound-specific gene expression of *Rpl13* and *Cidec* are shown in **Figure S4**.

### Adipogen Taxonomy Discovery

The taxonomy derived by the *K2Taxonomer* (Reed et al. 2020) procedure recapitulated many known characteristics shared by adipogens included in this study (**Figure 3**). For example, three terminal subgroups were labelled in **Figure 3** based on their shared characteristics. These include: flame retardants (tetrabromobisphenol A (TBBPA) and triphenyl phosphate (TPhP)), phthalates (MBUP, MEHP, MBZP, and BBZP), and RXR agonists (TBT and LG268).

**Figure 3.**
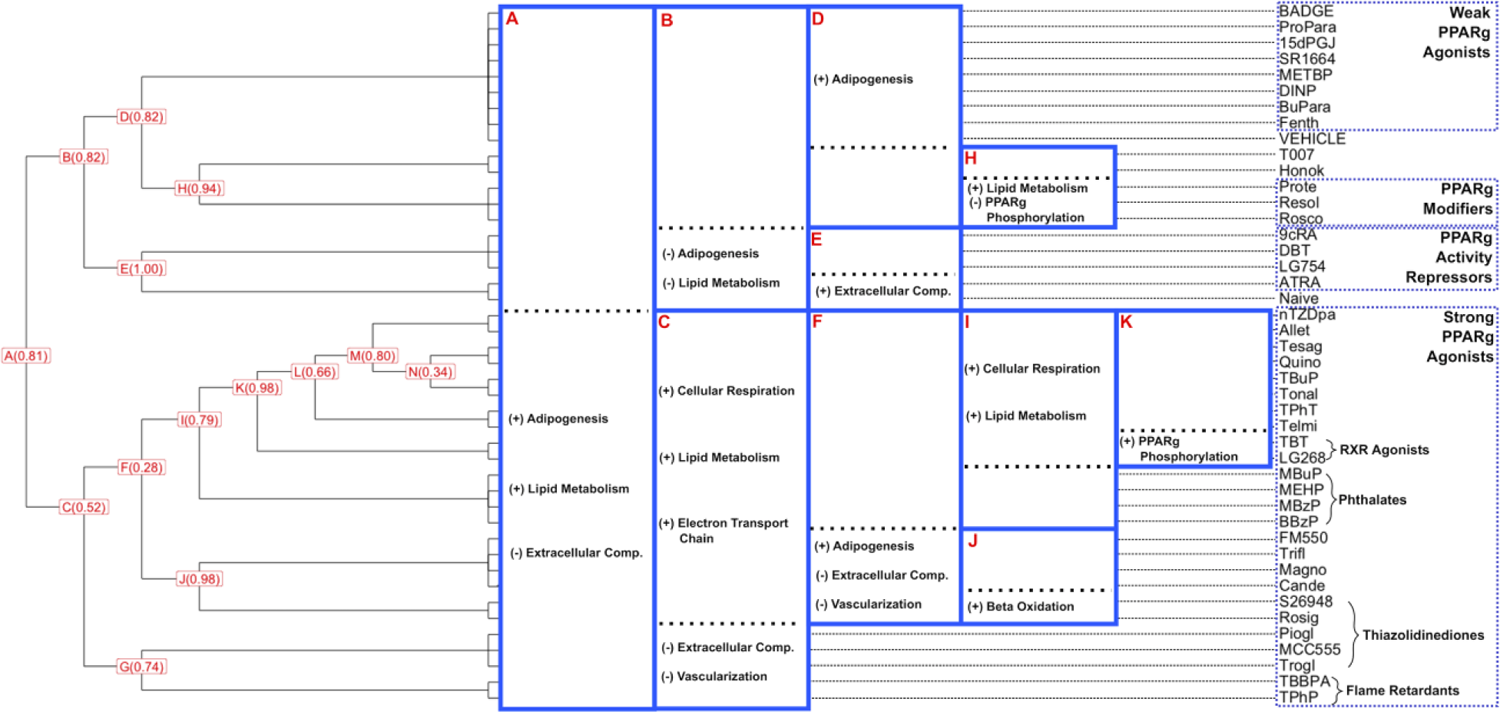
Chemical taxonomy of adipogens based on K2 clustering of the 3’DGE data. The dendrogram shows the taxonomy-driven hierarchical grouping of test chemical exposures of 3T3-L1 cells or naïve pre-adipocytes. Each split is labeled with a letter, and the proportion of gene-level bootstraps which produced the resulting split is shown. Highlights of hyper-enrichment of gene ontology (GO) biological processes are shown.

Interestingly, we observed two subgroups containing all of the four thiazolidinediones, with rosiglitazone (Rosig) segregating with the non-thiazolidinedione S26948 and pioglitazone, MCC 555, and troglitazone segregating together. Comprehensive functional annotation of the gene signatures and enriched pathways for comparisons between sub-groups at each split can be found in **Excel File 2**.

All of these terminal subgroups fell within a larger module containing 25 chemicals, highlighted by expression patterns consistent with higher adipogenic activity including up-regulation of genes significantly enriched in pathways involved in adipogenesis and lipid metabolism (Soukas et al. 2001). In addition, these chemicals also demonstrated consistent down-regulation of extracellular component genes. This effect was strongest in cells exposed to thiazolidinediones and flame retardants, two classes of chemicals well-described to be strong PPARγ agonists (Berger et al. 1996; Fang et al. 2015a; Riu et al. 2011). The subgroup of thiazolidinediones, including S26948, was characterized by up-regulation of genes involved in beta-oxidation, the process by which fatty acids are metabolized.

The gene expression profiles of the remaining 17 chemicals, including naïve pre-adipocytes, demonstrated markedly less up-regulation of genes regulated by PPARγ. Of these 17 perturbations, a subgroup of 8 chemicals (BADGE, PrPar, 15dPGJ2, SR1664, METBP, DINP, BuPA, and Fenth) included the reference vehicle signature. Compared to the next closest subgroup, expression profiles of these compounds were characterized by up-regulation of adipogenesis-related pathways indicative of modest PPARγ agonism. Additionally, a subgroup comprised of 9CRA, DBT, LG754, and ATRA exposures and naïve pre-adipocyte signatures was characterized by down-regulation of genes involved in adipogenesis and lipid metabolism, indicating repression of PPARγ activity. Interestingly, both protectin D1 (Prote) and resolvin E1 (Resol) clustered closely in a subgroup with the cyclin-dependent kinase (CDK) inhibitor, roscovitine (Rosco), which is known to induce insulin sensitivity and brite adipogenesis (Wang et al. 2016).

In summary, our top-down taxonomy discovery approach identified subgroups of adipogens characterized by differential transcriptomic activity at each split. Annotation of these transcriptomic signatures revealed clear differences in the set and magnitude of perturbations to known adipocyte biological processes by subgroups of chemicals. Membership of these subgroups confirmed many expectations, such as subgroups comprised solely of phthalates, thiazolidinediones, or flame retardants

### Relationship between taxonomic subgroups and the human adipose transcriptome

Given that our taxonomic clustering was based on adipogen exposures in a mouse model, we sought to establish its relevance to the relationship between human adipose tissue function and markers of cardio-metabolic health. To this end, we projected the gene signatures derived as either significantly increased or decreased gene sets in specific taxonomic subgroups onto a publicly available clinical data set of gene expression profiles from subcutaneous adipose tissue of 770 male subjects (Civelek et al. 2017) and assessed the relationship of these projections to a set of 12 clinical measurements. **Figure 4A-B** shows the partial correlation between plasma adiponectin and projection of gene sets, either increased or decreased in specific subgroups (further details are provided in **Excel File 4**). **Figure 4C** shows these relationships for the remaining measurements which demonstrated a statistically significant partial correlation for at least one projection, FDR Q-value < 0.10. For positive associations, i.e. in the same direction, the higher expressed gene sets had positive partial correlation measurements, while the lower expressed gene sets had negative partial correlation measurements. The opposite was true for negative associations.

**Figure 4.**
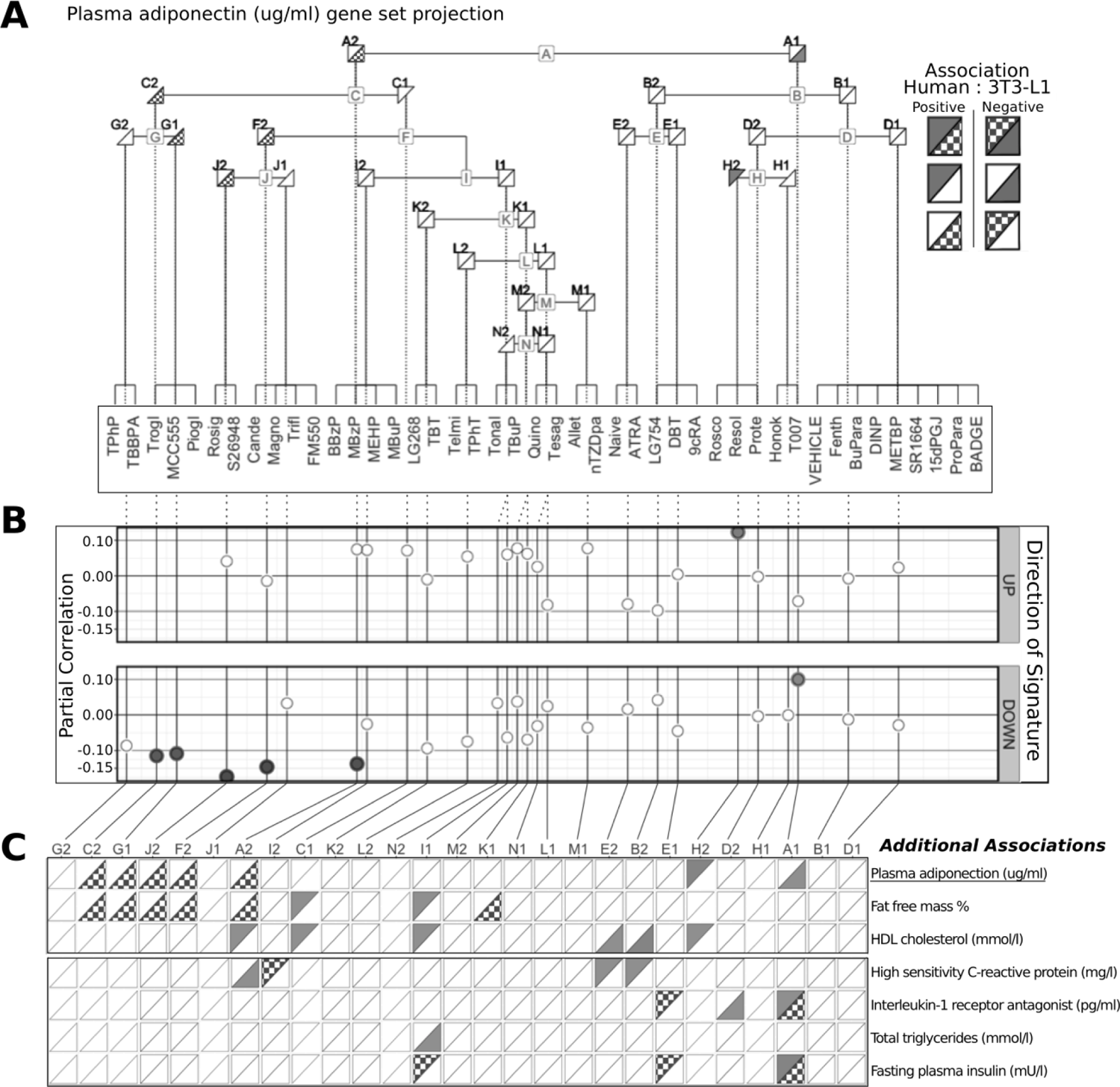
Associations between plasma adiponectin levels and projections of the 3T3-L1 derived chemical taxonomy gene signatures onto human adipose tissue gene expression. **(A)** Each adjoined triangle in the dendrogram denotes a set of genes derived from the 3T3-L1 data, either higher (left) or lower (right) expression at a given node. Triangle texture and color pertain to the direction of the partial correlation of the projection of these gene sets with plasma adiponectin levels, such that positive and negative partial correlations are indicated by solid red and checkered blue triangles, respectively. Interpretation of these associations is in the key on the upper right. All colored triangles reach a significance threshold of FDR Q-value < 0.10. **(B)** Plots of the partial correlation measurements for gene set projections and plasma adiponectin levels. The top and bottom plots are indicative of gene sets that are either increased in expression or decreased in expression in a particular sub-group, respectively. All colored points reach a significance threshold of FDR Q-value < 0.10, as in (A). **(C)** Results of all partial correlation analyses of projection of taxonomy gene sets with clinical measurements with at least one comparison reaching significance, FDR Q-value < 0.10. The interpretation of these associations is the same as in (A).

We observed concordant results for putative markers of metabolic health, plasma adiponectin and fat free mass %, which were positively associated with projection of gene signatures from two terminal subgroups which include all TZD and non-TZD type 2 diabetic drugs, Trogl, MCC555, Piogl, Rosig, and S26948, as well as a terminal subgroup which includes Rosco, Resol, and Protec. Similarly, we observed concordant results for putative markers of metabolic dysfunction, interleukin-1 receptor antagonist and fasting plasma insulin, which included a primary subgroup comprised of terminal subgroups characterized by weak PPARγ agonism, PPARγ modification, or PPARγ activity repression.

Taken together, these results confirm the ability of our mouse-based, in vitro-derived signatures of capturing salient functional aspects of healthy and unhealthy metabolic functions in human subjects.

### Investigation of the white and brite adipocyte taxonomy

We aimed to better assess how the distinction between gene expression patterns translated into functional differences in the induced adipocytes. Therefore, we selected chemicals from representative groups of related adipogens for genotypic and phenotypic characterization.

We compared a strong PPARγ therapeutic agonist that also modifies PPARγ phosphorylation (Rosig) (Choi et al., 2010), a chemical that modifies only PPARγ phosphorylation (Rosco) (Wang et al., 2016), a weak PPARγ agonist and endogenous molecule (15dPGJ2) (Foreman et al., 1995) and two known environmental PPARγ ligands (TBBPA and TPhP) (Chappell et al. 2018; Pillai et al. 2014). Protocols for differentiating 3T3 L1 cells are quite variable (reviewed and tested in Zhao et al. 2019), and there was some concern that the use of 250 nM dexamethasone would influence the differentiation process; therefore, we tested the effect of the ligands on gene expression in media with two dexamethasone concentrations (1 and 250 nM).

3T3-L1 cells were differentiated and treated with vehicle (DMSO), Rosig (positive control), Rosco, 15dPGJ2, TBBPA and TPhP. Cell density and gene expression were determined after 10 days. Rosig significantly increased cell number, while the other chemicals had no effect, positive or negative (**Figure S7**).

We determined if changes in gene expression correlated with expression of genes previously shown to be associated with white and brite/brown adipocytes (Qiang et al. 2012). As expected, Rosig, TBBPA, and TPhP significantly induced *Pparg2, Plin1, Fabp4* and *Cidec* expression in both dexamethasone conditions, while 15dPGJ2 only induced expression of these genes with 250 nM dexamethasone (**Figure 5A, Figure S8A**). *Pparg2* expression was moderately correlated between the two dexamethasone conditions (Spearman’s r = 0.06786, p = 0.0469), while *Plin1*, *Fabp4* and *Cidec* were highly correlated between conditions (Spearman’s r = 0.9286, p = 0.013; r = 0.8929, p = 0.0034; r = 0.9524, r < 0.0001, respectively). For genes expressed in brite adipocytes, a different pattern emerged, with little correlation between the two dexamethasone conditions. *Adipoq* is expressed by both white and brown adipocyte depots; it was induced by Rosig, TBBPA, and TPhP in both dexamethasone conditions. 15dPGJ2 and Rosco only induced *Adipoq* expression with 250 nM dexamethasone (**Figure 5A, Figure S8A**). For brite adipocyte genes, Rosig was the only ligand that induced expression of *Ppara*, *Pgc1a*, *Elovl3*, *Cidea*, *Ucp1*, *Acaa2* in both dexamethasone conditions (**Figure 5B**, **Figure S8B**). Rosco also induced expression of the brite adipocyte genes, but only in the 250 nM dexamethasone condition (**Figure S8B**). TBBPA and TPhP induced expression of *Ppara*, *Elovl3*, *Cidea*, *Acaa2* in low dexamethasone conditions only and did not induce expression of *Pgc1a* or *Ucp1* in either dexamethasone condition (**Figure 5B**, **Figure S8B**).

**Figure 5.**
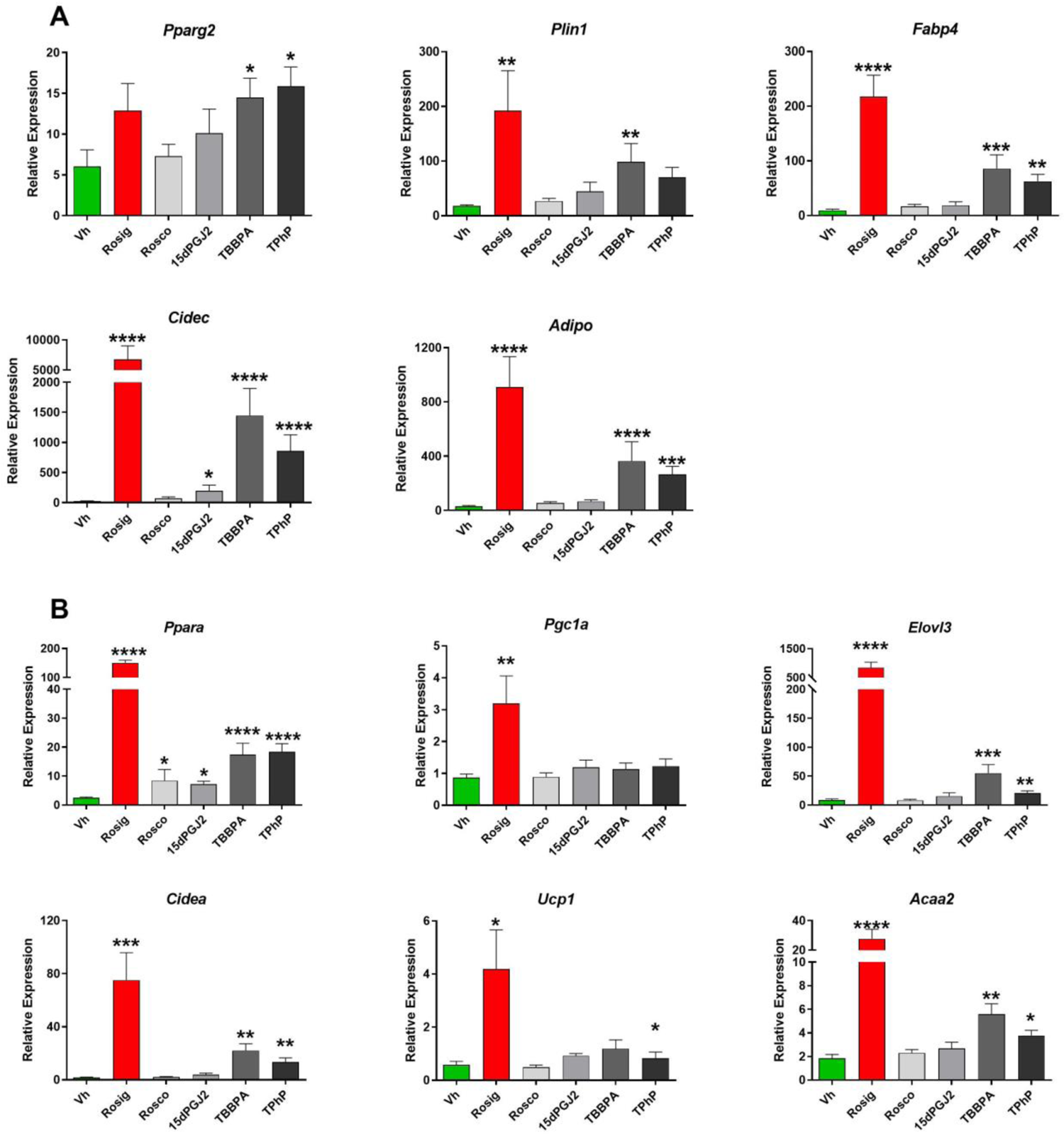
White and brite gene expression in differentiated and treated 3T3-L1 adipocytes. Confluent 3T3 L1 cells were differentiated using a standard hormone cocktail for 10 days. During differentiation, cells were treated with vehicle (Vh, 0.2% DMSO, final concentration), rosiglitazone (Rosig, 200 nM), roscovitine (Rosco, 4 μM), 15dPGJ2 (500 nM), TBBPA (20 μM) and TPhP (10 μM). On days 3, 5, and 7 of differentiation, the adipocyte maintenance medium was replaced and the cultures re-dosed. Following 10 days of differentiation and dosing, cells were analyzed for gene expression by RT-qPCR. Gene expression levels were normalized to the geometric mean of the expression levels of B2m and Rn18s and expressed as “Relative Expression” in comparison to naïve, pre-adipocyte cultures using the Pfaffl method. **(A)** Genes related to white adipogenesis. **(B)** Genes related to brite adipogenesis. Numerical data are provided in **Excel File 3**. Data are presented as mean <u>±</u> SE of n=4 independent experiments. Statistically different from Vh-treated (highlighted in green) (*p<0.05, **p<0.01, ***p<0.001, ****p<0.0001, ANOVA, Dunnett’s).

Next, we determined if changes in gene expression correlated with changes in adipocyte function. 3T3-L1 cells were differentiated in the presence of 1 nM dexamethansone and treated as described for the mRNA expression analyses. Compared to vehicle-treated cells, Rosig and TPhP significantly induced the rate of fatty acid uptake (**Figure 6A**) and Rosig, TPhP and TBBPA significantly increased total lipid accumulation (**Figure 6B**). Lipid accumulation was highly correlated with adiponectin secretion (**Figure 7**, Spearman’s r = 0.9565, p < 0.0001).

**Figure 6.**
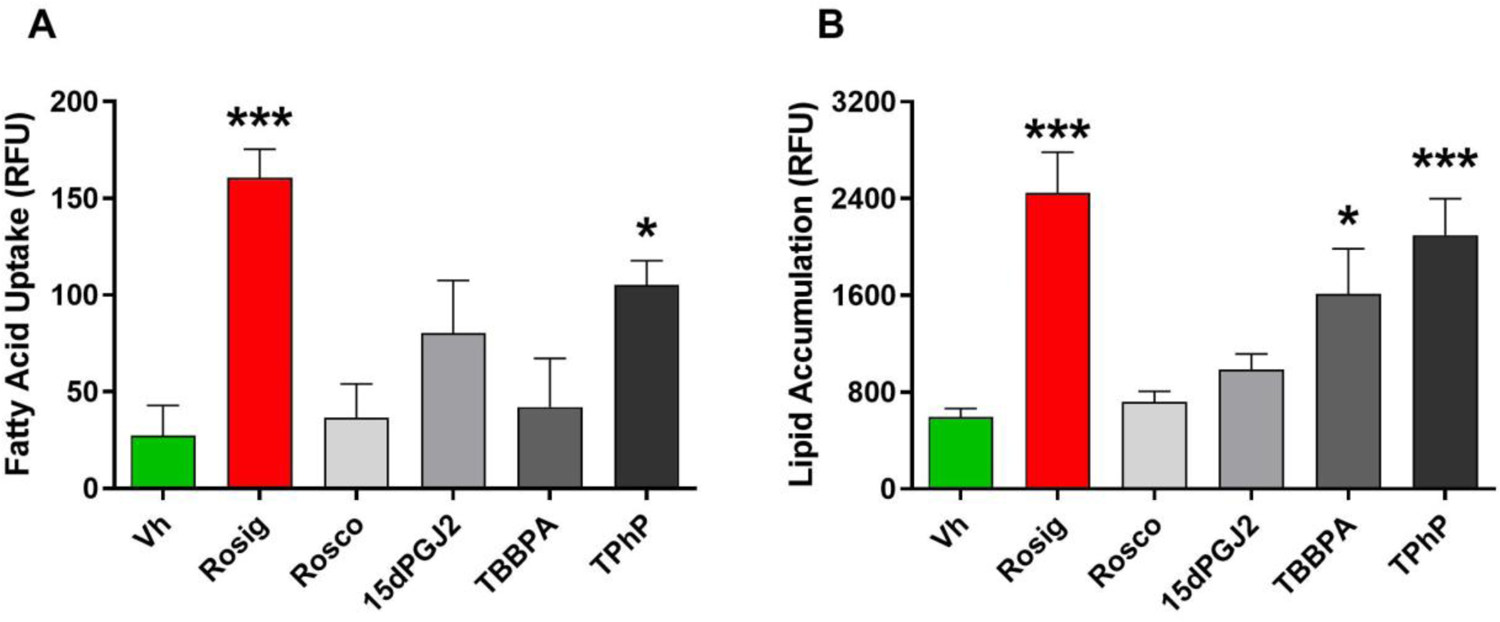
Fatty acid uptake and lipid accumulation in differentiated and treated 3T3-L1 adipocytes. Differentiation and dosing were carried out as described in Figure 5. Cells were analyzed after 10 days of differentiation. **(A)** Fatty acid uptake was analyzed using a dodecanoic acid fluorescent fatty acid substrate. The fluorescence at time zero was subtracted from the fluorescence at the end of the 10 min incubation and reported as “RFU.” **(B)** Lipid accumulation was analyzed by Nile Red staining. Nile Red fluorescence was normalized by subtracting the fluorescence measured in naïve pre-adipocyte cultures within each experiment and reported as “Naïve Corrected RFU.” Numerical data are provided in **Excel File 3**. Data are presented as means ± SE (n=4). Statistically different from Vh-treated (highlighted in green) (*p<0.05, ***p<0.001, ANOVA, Dunnett’s).

**Figure 7.**
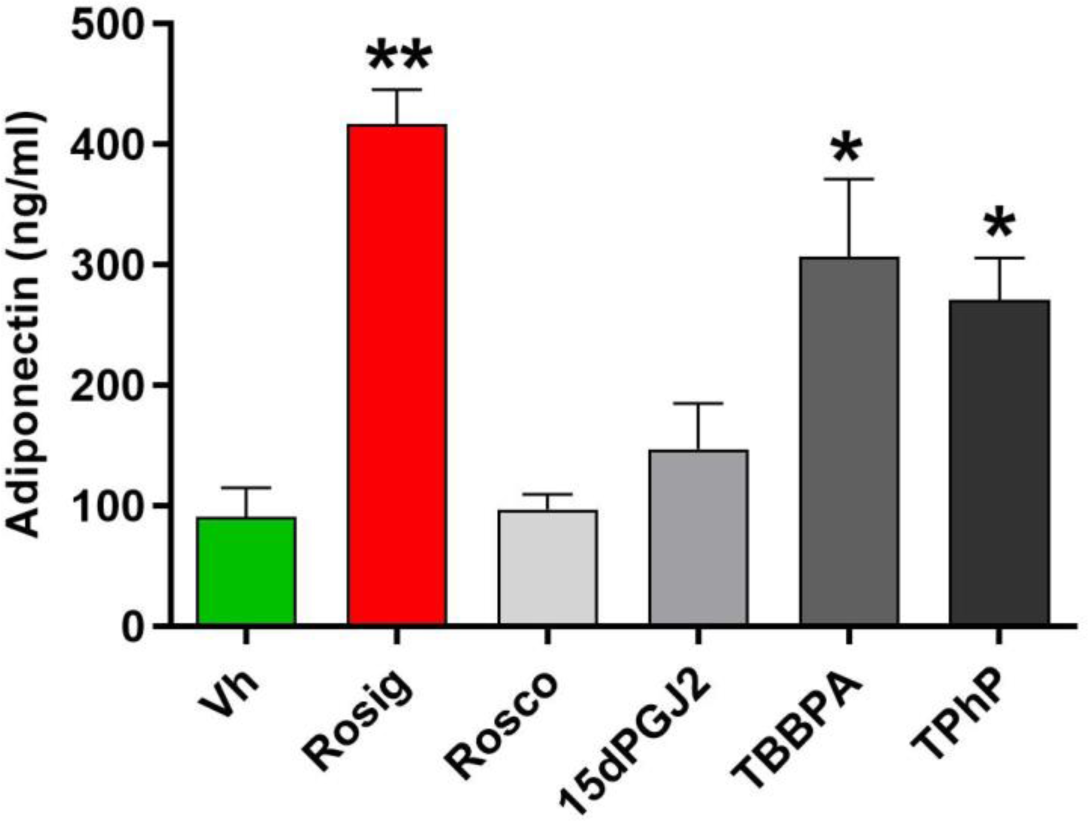
Adiponectin secretion in differentiated and treated 3T3-L1 adipocytes. Differentiation and dosing were carried out as described in Figure 5. On day 9 of differentiation, cells were switch to washout medium (DMEM+0.1% BSA) and incubated 24hrs. The washout medium was replaced and the cells were redosed. Medium was collected after a further 24hrs to adiponectin analysis by ELISA. Concentrations were calculated from absorbance values relative to a standard curve and reported as ng/ml. Numerical data are provided in **Excel File 3**. Data are presented as means ± SE (n=4). Statistically different from Vh-treated (highlighted in green) (*p<0.05, **p<0.01, ANOVA, Dunnett’s).

Overall, lipid accumulation was highly correlated with expression of all genes examined, with the exception of *Pgc1a* and *Ucp1* (**Figure S9**). Only Rosig significantly induced mitochondrial biogenesis (**Figure 8**). Rosig also significantly increased cellular respiration (**Figure 9**). Rosco, 15dPGJ2, TBBPA and TPhP did not change cellular respiration.

**Figure 8.**
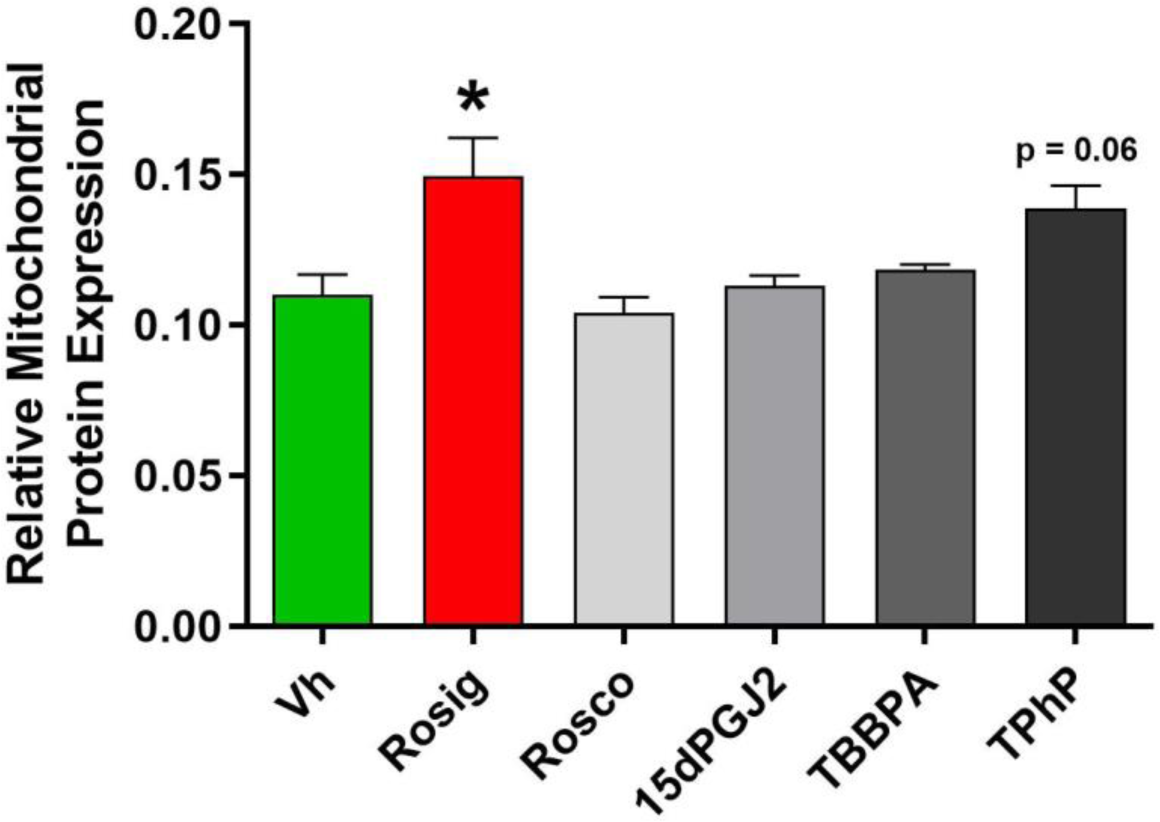
Mitochondrial biogenesis in differentiated and treated 3T3-L1 adipocytes. Differentiation and dosing were carried out as described in Figure 5. Following 10 days of differentiation, mitochondrial biogenesis was analyzed by measuring mitochondria-specific proteins by ELISA. The absorbance ratios of SDH/Janus are reported as “Relative Mitochondrial Protein Expression.” Numerical data are provided in **Excel File 3**. Data are presented as means ± SE (n=4). Statistically different from Vh-treated (highlighted in green) (*p<0.05, ANOVA, Dunnett’s).

**Figure 9.**
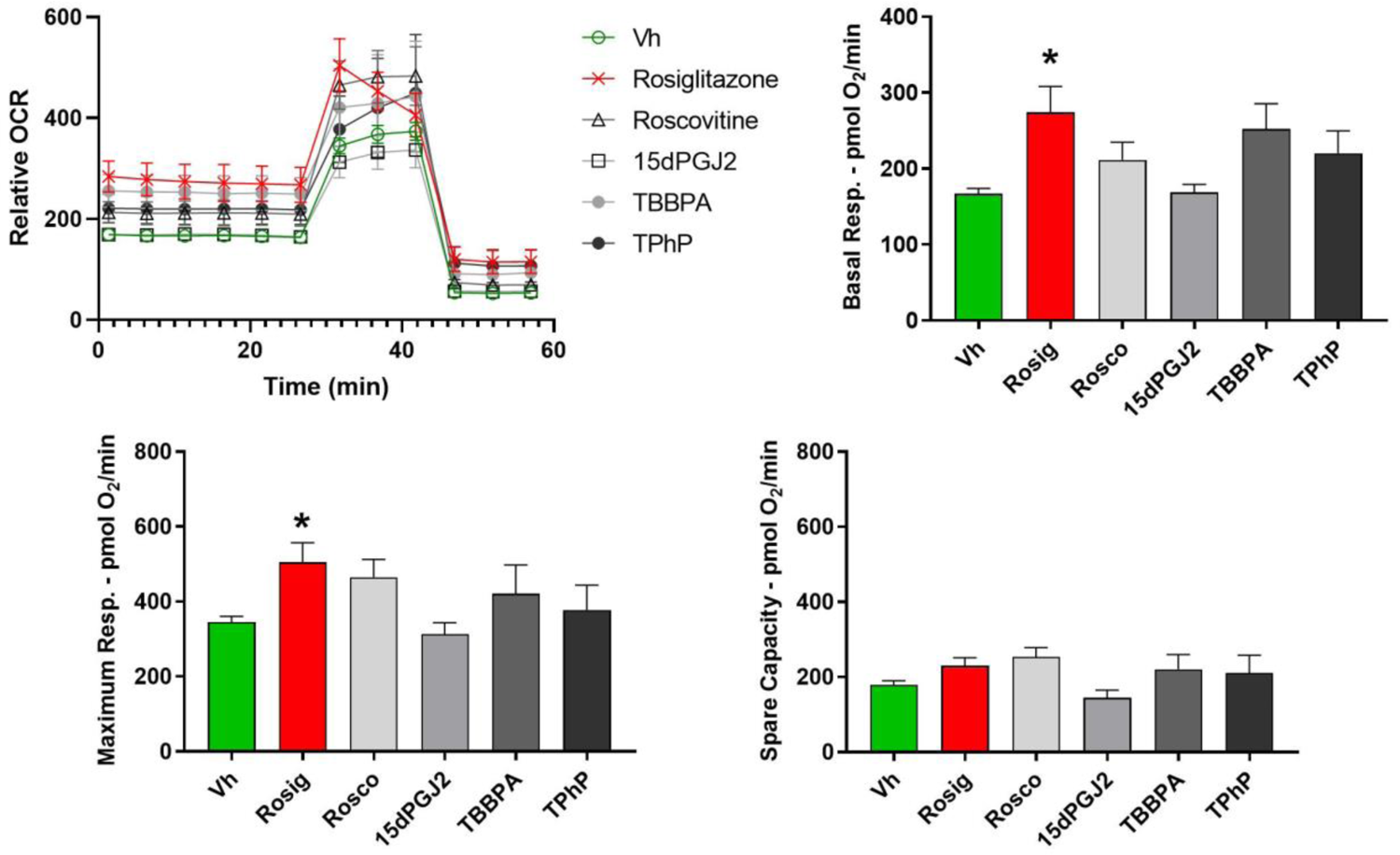
Cellular respiration in differentiated and treated 3T3-L1 adipocytes. Differentiation and dosing were carried out as described in Figure 5. Following 10 days of differentiation, mitochondrial respiration was analyzed by Seahorse Assay. Oxygen consumption rate was normalized for cell number by dividing by the Janus Green absorbance in each well and reported as “Relative OCR.” Numerical data are provided in **Excel File 3**. Data are presented as means ± SE (n=4). Statistically different from Vh-treated (highlighted in green) (*p<0.05, ANOVA, Dunnett’s).

### Identification of adipogens that favor white adipogenesis

Quinoxyfen (Quino) and tonalide (Tonal) were two of the environmental chemicals that received the highest PPARγ ligand/modifier vote and segregated distinctly from the therapeutic ligands (**Table 2**). Thus, we tested the hypothesis that Quino and Tonal are adipogens that do not induce gene expression or metabolic phenotypes indicative of high energy expenditure or brite adipogenesis. 3T3-L1 cells were differentiated in the presence of 1 nM dexamethasone and treated with vehicle (DMSO), rosiglitazone (positive control), Quino, or Tonal. These concentrations were determined to be non-toxic with only Rosig increasing cell number (**Figure S10A**). In 3T3-L1 cells, Rosig and Quino significantly increased the rate of fatty acid uptake, but Rosig, Quino and Tonal all significantly increased total lipid accumulation (**Figure 10A**). Rosig and Quino, but not Tonal, significantly increased adiponectin secretion (**Figure 10B**).

Interestingly, only Quino significantly increased the expression of *Pparg2.* Both Rosig and Quino significantly increased expression of *Plin1*, *Cidec*, and *Cidea*, while Tonal had little efficacy in inducing expression of adipocyte-specific genes (**Figure 10C**). While both Rosig and Quino significantly upregulated expression of *Cidea*, only Rosig significantly induced *Pgc1a* and *Ucp1* (**Figure 10D**). In spite of both Rosig and Quino both increasing mitochondrial biogenesis (**Figure 10E**), only Rosig significantly increased cellular respiration (**Figure 10F**).

**Figure 10.**
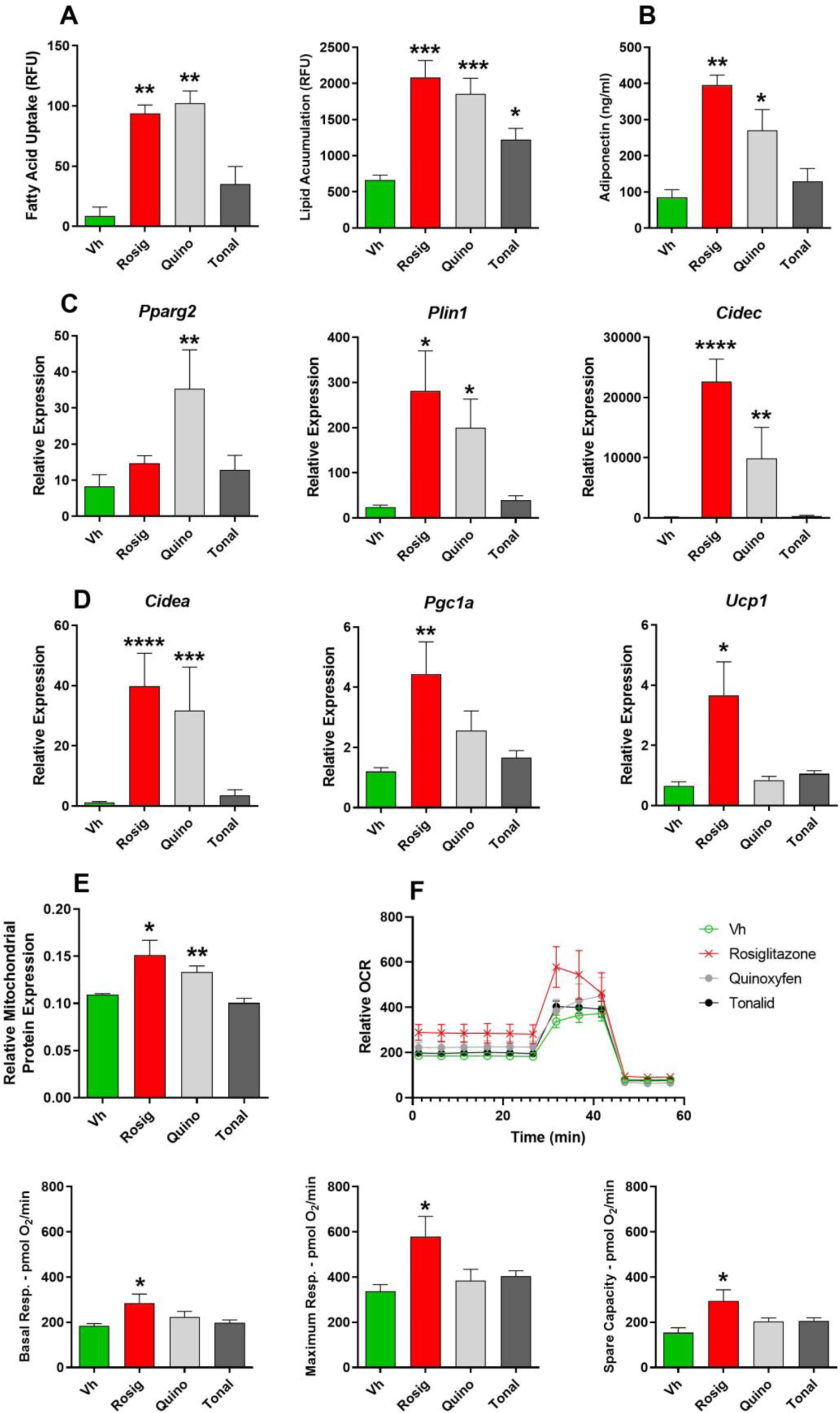
Quinoxyfen- and tonalide-induced adipogenesis in 3T3-L1 pre-adipocytes. Confluent 3T3 L1 cells were differentiated using a standard hormone cocktail for 10 days. During differentiation, cells were treated with vehicle (Vh, 0.2% DMSO, final concentration), rosiglitazone (Rosig, 200 nM), quinoxyfen (Quino, 10 μM) or tonalide (Tonal, 4 μM). On days 3, 5, and 7 of differentiation, the adipocyte maintenance medium was replaced and the cultures re-dosed. Following 10 days of differentiation and dosing, cultures were analyzed. **(A)** Lipid handling. For fatty acid uptake, the fluorescence at time zero was subtracted from the fluorescence at the end of the 10 min incubation and reported as “RFU.” Lipid accumulation was determined from Nile Red fluorescence, which was normalized by subtracting the fluorescence measured in naïve pre-adipocyte cultures within each experiment and reported as “Naïve Corrected RFU.” **(B)** Adiponectin concentrations in the medium measured by ELISA and calculated from absorbance values relative to a standard curve and reported as ng/ml. Gene expression levels determined by RT-qPCR and were normalized to the geometric mean of the expression levels of *B2m* and *Rn18s* and expressed as “Relative Expression” in comparison to naïve, pre-adipocyte cultures using the Pfaffl method. **(C)** White adipocyte gene expression. **(D)** Brite adipocyte gene expression. **(E)** Mitochondrial biogenesis was determined by measuring protein concentrations by ELISA. Absorbance ratios of SDH/Janus are reported as “Relative Mitochondrial Protein Expression.” **(F)** Cellular respiration was measured by Seahorse Assay. Oxygen consumption rate was normalized for cell number by dividing by the Janus Green absorbance in each well and reported as “Relative OCR.” Numerical data are provided in **Excel File 3**. Data are presented as means ± SE (n=4-6). Statistically different from Vh-treated (highlighted in green) (*p<0.05, **p<0.01, ***p<0.001, ANOVA, Dunnett’s).

Last, we investigated whether results in our mouse model, 3T3-L1 cells, could be recapitulated in a human model. Primary, human subcutaneous preadipocytes were differentiated and treated with vehicle (DMSO), rosiglitazone (positive control), Quino, or Tonal. Quino and Tonal significantly induced lipid accumulation (**Figure 11A**). Tonal, but not Quino, increased cell number (**Figure S10B**). Both Quino and Tonal induced *CIDEC* expression but failed to induce *CIDEA* expression (**Figure 11B**). In contrast to 3T3-L1 cells, Quino and Tonal did not increase fatty acid uptake over that induced by the hormonal cocktail in the differentiated primary human adipocytes (**Figure 11C**). Quino and Tonal did not induce mitochondrial biogenesis (**Figure 11D**).

**Figure 11.**
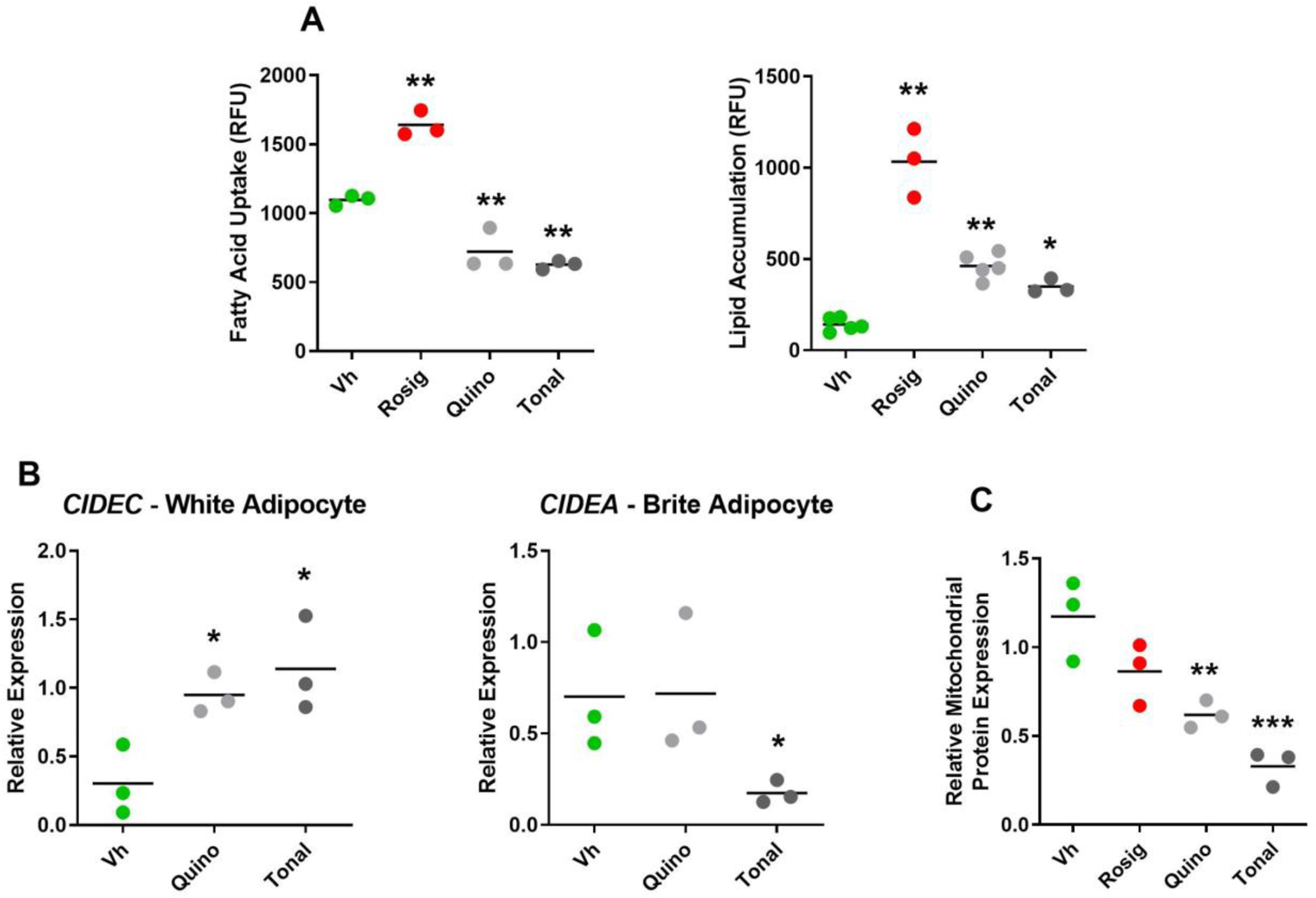
Tonalide- and quinoxyfen-induced adipogenesis in primary human adipocytes. Confluent primary human preadipocytes were differentiated using a standard human adipocyte hormone cocktail for 14 days. During differentiation, cells were treated with vehicle (Vh, 0.1% DMSO, final concentration), rosiglitazone (Rosig, 4 μM), quinoxyfen (Quino, 4 μM) or tonalide (4 μM). On days 3, 5, 7, 10, and 12 of differentiation, the medium was replaced and the cultures re-dosed. Following 14 days of differentiation and dosing, cultures were analyzed for **(A)** lipid handling, **(B)** white (*Cidec*) and brite (*Cidea*) gene expression, and **(C)** mitochondrial biogenesis. Individual data are presented with the mean indicated by a line. Gene expression levels were normalized to the geometric mean of the expression levels of *B2M* and *RPL27* and expressed as relative expression in comparison to naïve, pre-adipocyte cultures. Numerical data are provided in **Excel File 3**. Statistically different from Vh-treated (highlighted in green) (*p<0.05, **p<0.01, ***p<0.001, ANOVA, Dunnett’s).

## Discussion

The chemical environment has changed dramatically in the past 40 years, and an epidemic increase in the prevalence of obesity has occurred over the same time period (Baillie-Hamilton 2002). Yet, it is still unclear how chemical exposures may be contributing to adverse metabolic health effects. New tools are needed not just to identify potential adipogens, but to provide information on the type of adipocyte that is formed. Here, we have both developed a new analytical framework for adipogen identification and characterization and tested its utility in hypothesis generation. We showed that adipogens segregate based on distinct patterns of gene expression. Furthermore, subgroups within the taxonomy containing therapeutics shared gene expression patterns in human adipose tissue in accordance with positive markers of metabolic health, specifically plasma adiponectin and fat free mass %, suggesting that these gene expression are related to metabolic health in humans. We identified two environmental contaminants segregating outside the therapeutic groups for hypothesis testing. Our results support the conclusion that quinoxyfen and tonalide have a limited capacity to induce the health-promoting effects of mitochondrial biogenesis and brite adipocyte differentiation.

### Adipogen taxonomy identifies environmental chemicals that favor white adipogenesis

Potential adipogens (chemicals that change the differentiation and/or function of adipocytes) were identified by review of the literature and based on reports of PPARγ agonism or modulation of adipocyte differentiation, as well as through querying ToxCast data. Not all of the chemicals previously identified as adipogens induced significant lipid accumulation.

Mono(2-ethylhexyl) phthalate (MEHP)(Feige et al. 2007), SR1664 (Choi et al. 2011), and 15-deoxy-Δ^12,14^-prostaglandin J2 (15dPGJ2)(Forman et al. 1995) are PPARγ agonists that were expected to increase adipocyte differentiation but did not. LG268 (Boehm et al. 1994) and TBT are RXR agonists that were also expected to significantly increase adipocyte differentiation but did not. T007 is a PPARγ antagonist (Lee et al. 2002) that was expected to decrease adipocyte differentiation but did not. We hypothesize that this likely resulted from the fact that we did not apply any chemical above 20 μM (with the exception of fenthion). Concentrations in the 100 μM range have been used in previous studies (e.g. DOSS (Temkin et al. 2016); parabens (Hu et al. 2013); phthalates (Hurst and Waxman 2003; Feige et al. 2007)). The RXR agonists LG268 and TBT also were expected to significantly increase adipocyte differentiation but did not.

We tested the chemicals in multiple differentiation scenarios (3T3-L1s with 0, 1 or 250 nM dexamethasone and in OP9 cells). Each differentiation scenario generated highly correlated results, but with some deviations. 3T3-L1 cells are the most used model in the study of adipogens (Green and Kehinde 1975; Ruiz-Ojeda et al. 2016), with a broad range of “standard” differentiation cocktails. Not surprisingly, in 3T3 L1 cells differentiated in the absence of dexamethasone, glucocorticoid receptor agonists significantly induced lipid accumulation. A significant drawback of this model, however, is the variability of the efficacy of differentiation, which has been shown to depend on the commercial source, passage number, culture dishes used and medium volume used (Sheng et al. 2014; Kassotis et al. 2017). On the other hand, an important feature of 3T3-L1 cells is their ability to differentiate into both white and brite adipocytes (Vernochet et al. 2009). Thus, we suggest that the minimum amount of dexamethasone to support efficient differentiation, without inducing significant background differentiation, should be identified for a given lot of cells before proceeding with testing.

We have previously shown that TBT induced adipogenesis with greater efficacy in OP9 cells than in 3T3-L1 cells (Kassotis et al. 2017; Andrews et al., 2020), and also showed here that OP9 cells more efficaciously responded to RXR ligands, in general, likely because they are more committed to adipogenesis as late pre-adipocytes (Wolins et al., 2006) than 3T3 L1 cells, which are early pre-adipocytes (Lane and Tang, 2005). OP9 are significantly less finicky than 3T3-L1 cells in terms of the experimental conditions required for efficient adipocyte differentiation, likely because they are further along in commitment to adipogenesis than 3T3-L1 cells (Wolins et al., 2006; Kassotis et al., 2017). However, they have not been tested for their ability to differentiate into brite adipocytes.

We identified four compounds as high-confidence PPARγ ligands/modifiers: quinoxyfen, tonalide, allethrin, and fenthion. In the final model used for these predictions, biologically informative genes emerged as important for predicting PPARγ ligand/modification status, specifically the down-regulation of *Rpl13* and the up-regulation of *Cidec. Rpl13* is involved in ribosomal machinery and is down-regulated during human adipogensis (Marcon et al. 2017).

*Cidec* is a lipid droplet structural gene, the expression of which was positively correlated with adipocyte lipid droplet size, insulin levels, and glycerol release (Ito et al. 2010). Of these four compounds, quinoxyfen and tonalide are of particular public health concern. Quinoxyfen is among a panel of pesticides with different chemical structures and modes of action (i.e., zoxamide, spirodiclofen, fludioxonil, tebupirimfos, forchlorfenuron, flusilazole, acetamaprid, and pymetrozine) that induced adipogenesis and adipogenic gene expression in 3T3-L1 cells (Janesick et al. 2016). Quinoxyfen is a fungicide widely used to prevent the growth of powdery mildew on grapes (Duncan et al. 2018). We chose to test tonalide because it was reported to strongly increase adipogenesis in 3T3-L1 cells, although it was concluded that this response was not due to direct PPARγ activation (Pereira-Fernandes et al. 2013). Our results differed in this regard. Tonalide bioaccumulates in adipose tissue of many organisms including humans, and exposure is widespread because of its common use in cosmetics and cleaning agents (Kannan et al. 2005). Combined, tonalide and galaxolide constitute 95% of the polycyclic musks used in the EU market and 90% of that of the US market (HERA 2004).

Our results support the conclusion that quinoxyfen and tonalide are adipogenic chemicals, likely to be acting through PPARγ. In clustering analysis, quinoxyfen and tonalide were among the largest subgroup of eight potential strong PPARγ agonists. Notably, this cluster included both synthetic/therapeutic (nTZDpa, tesaglitazar, telmisartan) and environmental compounds (tributyl phosphate and triphenyltin) and was characterized by general up-regulation of pathways of adipogenic activity. However, quinoxyfen and tonalide generated adipocytes that were phenotypically distinct from adipocytes induced by therapeutics such as rosiglitazone. That environmental PPARγ ligands can induce a distinct adipocyte phenotype has been shown previously for TBT (Kim et al. 2018; Regnier et al. 2015; Shoucri et al. 2018) and TPhP (Kim et al. 2020). Ultimately, binding assays or computational ligand binding domain modeling will be needed to confirm that quinoxyfen and tonalide are PPARγ agonists.

We tested the effect of quinoxyfen and tonalide on human subcutaneous preadipocyte differentiation. There was a strong association between abdominal adiposity and metabolic syndrome (Cornier et al. 2008). Classically, visceral adipose tissue has been thought to be the driver of metabolic dysfunction; however, there is an alternative explanation that visceral adiposity results secondarily from the dysfunction of subcutaneous adipose tissue in the upper body (Jensen 2008; Lee et al. 2017). Subcutaneous adipose tissue represents 85% of all body fat (Frayn and Karpe 2014) and thus has a large overall capacity for generating brite adipocytes.

Additionally, lack of browning capacity of human subcutaneous adipocytes was associated with insulin resistance in a group of male Swedish individuals (Yang et al. 2003). As hypothesized based on the taxonomical analysis, quinoxyfen and tonalide induced white adipocyte functions such as increased lipid accumulation, but in contrast to rosiglitazone, did not induce mitochondrial biogenesis.

We hypothesize that the differences in adipocyte phenotype that are induced by environmental adipogens and likely PPARγ ligands (e.g., TBBPA, TPhP, quinoxyfen, tonalide) could result from the conformation that PPARγ assumes when liganded with these chemicals rather than with therapeutic agents. Differences in conformation not only determine the efficacy to which PPARγ is activated but also the transcriptional repertoire (Chrisman et al. 2018). We have shown recently, for instance, that TPhP did not protect PPARγ from phosphorylation at s273 and did not induce brite adipogenesis; however, when PPARγ cannot be phosphorylated at s273, TPhP could induce brite adipogens (Kim et al. 2020). Structural analyses would need to be conducted with each environmental ligand specifically to determine the contribution of conformational changes to the biological activity.

Access to post-translational modification sites and coregulator binding surfaces depends upon the structure that PPARγ assumes. The white adipogenic, brite/brown adipogenic and insulin sensitizing activities of PPARγ are regulated separately through differential co-regulator recruitment (Villanueva et al. 2013) and post-translational modifications (Choi et al. 2010; Choi et al. 2011), with ligands having distinct abilities to activate each of PPARγ’s functions. Suites of genes have been shown to be specifically regulated by the acetylation status of PPARγ (SirT1-mediated) (Qiang et al. 2012), by the phosphorylation status of PPARγ (ERK/MEK/CDK5-mediated) (Choi et al. 2010; Wang et al. 2016) and/or by the recruitment of Prdm16 to PPARγ (Seale et al. 2007). Future work will investigate the connections between the phosphorylation status of PPARγ liganded with environmental PPARγ ligands, the recruitment and release of coregulators, and the ability of PPARγ to recruit transcriptional machinery to specific DNA-binding sites. It will be important to determine the metabolic effects of chemicals like TPhP, quinoxyfen and tonalide *in vivo*.

### Analytical approaches for adipogen characterization

In this study, we performed a high-throughput, cost-effective transcriptomic screening to profile adipocytes formed from 3T3-L1 preadipocytes exposed to a panel of compounds of known and unknown adipogenic impact. Common to toxicogenomic projects, this panel-based study design allowed for characterization of the extent to which each chemical modified differentiation (in this case, adipogenesis as related to the change in lipid accumulation). It also supported the exploration of how subsets of chemicals influence multiple biological processes that determine the functional status of a cell (in this case, processes that determine white *vs.* brite adipogenesis). Exploration of these biological processes allowed for the prediction of the phenotypic impact of previously unclassified compounds, as well as for the characterization of the heterogeneity of the cellular activity of compounds with similar known phenotypic impact.

Here, we performed both types of analyses: first through the implementation and application of random forest classification models to identify potential adipogens, and second via the recursive clustering of the data to identify and characterize taxonomic subgroups of known and potential PPARγ ligands/modifiers.

For both analyses, we introduced amendments to commonly used machine learning procedures, to improve accuracy and resolution of the acquired result. For the classification task, we amended the random forest algorithm to tailor it to study designs typically adopted in toxicogenomic projects (see Methods). With the addition of an extra step to average the expression across replicates of the bootstrapped samples, we observed consistently higher performance across conventional metrics than with the standard algorithm. For the clustering task, we employed a procedure where we recursively divide sets of chemicals into two subgroups and assessed the robustness of each division, as well as annotated transcriptional drivers of each division. As a result, we were not limited to interpreting the clustering results as mutually exclusive groups, but rather as a taxonomy of subgroups where sets of compounds shared some transcriptional impact and differ in others, as was expected given the dynamic nature of the modifications by which compounds directly and indirectly affect PPARγ activity.

Future work will generalize random forest method to incorporate more complex study designs. To this end, the classification approach adopted in this project is being developed as a random forest software tool soon to be made available as an R package, allowing for the interchanging independent functions at different steps of the algorithm. The strength and utility of this approach extends beyond toxicogenomic studies, and can be used in a variety of applications of high-throughput screening, including drug discovery, such as the Connectivity Map (CMAP) (Subramanian et al. 2017), and longitudinal molecular epidemiology studies, such as the Framingham Heart Study (Mahmood et al. 2014).

### Adipogen portal

Given the breadth of results generated by this analysis, our description here is far from exhaustive. As such, we have created an interactive website (https://montilab.bu.edu/adipogenome/) to support the interactive exploration of these results at both the gene and pathway-level. A help page illustrating how to navigate this web portal is provided on the website as well. The portal is built around a point-and-click dendrogram of the clustering results as in **Figure 3**. Selecting a node of this dendrogram will populate the rest of the portal with the chemical lists, differential analysis, and pathway level hyper-enrichment results for each subgroup defined by a split. For instance, selecting node “H” will show the chemicals in each subgroup to the right (Group 1 = Honokiol, T007907; Group 2 = Prote, Resol, and Rosco), as well as the differential gene signature for each group below. Selecting *Cidec*, the top gene in the Group 2 signature, displays hyper-enrichment results for gene sets which include *Cidec* and have a nominal p-value < 0.50. The hyper-enrichment results for all genes can be found below this table. Finally, selecting a gene set name will display the gene set members at the bottom frame of the portal, with gene hits in bold. All tables are query-able and downloadable.

## Conclusions

Emerging data implicate contributions of environmental metabolism-disrupting chemicals to perturbations of pathways related to metabolic disease pathogenesis, such as disruptions in insulin signaling and mitochondrial activity. There is still a gap in identifying and examining how environmental chemicals can act as obesity-inducing and metabolism-disrupting chemicals. Our implementation of novel strategies for classification and taxonomy development can help identify environmental chemicals that are acting on PPARγ. Further, our approach provides a basis from which to investigate effects of adipogens on not just the generation of adipocytes, but potentially pathological changes in their function. To this end, we have shown how two environmental contaminants, quinoxyfen and tonalide, are inducers of white adipogenesis. Finally, we emphasize that these approaches are generally suitable for analyses of transcriptomic data generated from toxicogenomic screening studies. For example, provided such data, *K2Taxonomer* (Reed et al. 2020) can be applied to characterize molecular patterns driving segregation of experimental groups and infer dysregulated biological activity inherent to subgroups of experimental groups, thereby providing insight to inform Adverse Outcome Pathway (AOP) models of response to chemical exposures (Brockmeier et al. 2017).

## Notes

### Competing Interest Statement

The authors have declared no competing interest.

### Summary of Updates

New data appear in Figures 5-10.

