## Supplemental Material for "A Data-Driven Transcriptional Taxonomy of Adipogenic Chemicals to Identify White and Brite Adipogens"

### Table of Contents


| | | Page |
|---|---|---|
| **Table S1** | Chemical information. | **2** |
| **Table S2** | Mouse and human primer sequences for reverse transcriptase qPCR. | **7** |
| **Table S3** | Metabolic parameters included and excluded in human transcriptome analysis. | **7** |
| **Excel File 1** | Detailed results of final random forest model. | **8** |
| **Excel File 2** | Detailed annotation of clustering results for individual modules. | **8** |
| **Excel File 3** | Supporting numerical data from all cell culture experiments. | **8** |
| **Excel File 4** | Detailed results of partial correlation analysis of clinical measurements and projections of chemical taxonomy gene signatures onto human adipose gene expression. | **8** |
| **Figure S1** | Differentiation protocols | **9** |
| **Figure S2** | Correlation of lipid accumulation with *Cidec* expression in differentiated and treated 3T3-L1 pre-adipocytes. | **10** |
| **Figure S3** | Lipid accumulation in differentiated and treated 3T3-L1 pre-adipocytes in the absence of dexamethasone. | **11** |
| **Figure S4** | Lipid accumulation in differentiated and treated OP9 pre-adipocytes. | **13** |
| **Figure S5** | Performance comparison of random forest methods. | **15** |
| **Figure S6** | Classification results (distributions of individual genes) | **16** |
| **Figure S7** | Cell number analyses in the differentiated and treated 3T3-L1s. | **18** |
| **Figure S8** | White and brite gene expression in differentiated and treated 3T3-L1 adipocytes differentiated in 250 nM dexamethasone. | **19** |
| **Figure S9** | Spearman correlation analysis of lipid accumulation (Nile Red) and gene expression. | **21** |
| **Figure S10** | Cell number analyses in the differentiated and quinoxyfen and tonalide treated 3T3-L1s and human primary preadipocytes. | **22** |
| | **References** | **19** |

**Table S1. Chemical information.**

| Chemical Name | Abbr. | CAS # | Supplier | Catalog # | Max. Conc. Tested [M] | Max. Non-Toxic Conc. [M][a] | PPARγ Ligand or Modifier[b] | Purity | Reference |
|---|---|---|---|---|---|---|---|---|---|
| **15-deoxy-Δ12,14-prostaglandin J2** | 15dPGJ | 87893-55-8 | Cayman Chemical | 18570 | 1x10-6 | 1x10-6 | Yes PPARγ ligand | > 95% | (Forman et al. 1995) |
| **2,2',4,4',5,5'-Hexachloro-1,1'-biphenyl** | PCB153 | 35065-27-1 | Ultra Scientific | RPC-047 | 1x10-5 | 1x10-5 | No evidence | NA* | --- |
| **2,2',5,5'-Tetrachloro-1,1'-biphenyl** | PCB52 | 35693-99-3 | Sigma Aldrich | 35599 | 1x10-5 | 1x10-5 | No evidence | > 98% | --- |
| **2,4,6-Tris(tert-butyl)phenol** | TTBP | 732-26-3 | Sigma Aldrich | T49409 | 2x10-5 | 2x10-5 | Potential | 98% | (Auerbach et al. 2016) |
| **2-ethylhexanol** | EtHex | 104-76-7 | Sigma Aldrich | W315109 | 1x10-5 | 1x10-5 | No evidence | > 99% | --- |
| **3,3',4,4',5-Pentachloro-1,1'-biphenyl** | PCB126 | 57465-28-8 | Ultra Scientific | RPC-102 | 1x10-8 | 1x10-8 | No AhR[c] ligand, Reduces adipogenesis | NA | (Gadupudi et al. 2015) |
| **3,3',5,5'-Tetrabromobisphenol A** | TBBPA | 79-94-7 | Sigma Aldrich | 330396 | 2x10-5 | 2x10-5 | Yes PPARγ ligand | 97% | (Riu et al. 2011) |
| **4,4'-Dichloro-diphenyldichloroethylene** | DDE | 72-55-9 | Sigma Aldrich | 48679 | 1x10-5 | 1x10-5 | No ER ligand | NA | (Kim et al. 2016) |
| **4,4'-dichloro-diphenyltrichloroethane** | DDT | 50-29-3 | Sigma Aldrich | 40124 | 1x10-5 | 1x10-5 | No ER ligand | NA | (Kim et al. 2016) |
| **4,5,6,7-Tetrabromobenzotriazole** | TBB | Synthesized by Asis Chemical | | | 1x10-5 | 1x10-5 | No evidence | 95% | --- |
| **9-cis-retinoic acid** | 9cRA | 5300-03-8 | Sigma Aldrich | R4643 | 1x10-6 | 1x10-6 | Yes Activates PPARγ through RXR | > 98% | (Szeles et al. 2010) |
| **All-trans retinoic acid** | ATRA | 302-79-4 | Sigma Aldrich | R2625 | 2x10-6 | 2x10-6 | No RAR ligand, Reduces adipogenesis | > 98% | (Schwarz et al. 1997) |
| **Benzyl butyl phthalate** | BBzP | 85-68-7 | Sigma Aldrich | 36927 | 1x10-5 | 1x10-5 | Yes Induces PPARγ target genes and 3T3 L1 adipogenesis | 98% | (Yin et al. 2016) |
| **Bisphenol A** | BPA | 80-05-7 | Sigma Aldrich | 239658 | 1x10-5 | 1x10-5 | No ER ligand | > 99% | (Molina-Molina et al., 2013) |
| **Bisphenol A diglycidyl ether** | BADGE | 1675-54-3 | Sigma Aldrich | D3415 | 1x10-5 | 1x10-5 | Yes Induces PPARγ target genes and | NA | (Chamorro-Garcia et al. 2012) |

| | | | | | | | | | |
|---|---|---|---|---|---|---|---|---|---|
| | | | | | | | 3T3 L1 and stromal cell adipogenesis | | |
| **Bisphenol S** | BPS | 80-09-1 | Sigma Aldrich | 43034 | 1x10-5 | 1x10-5 | No<br>ER ligand | NA | (Molina-Molina et al., 2013) |
| **Candesartan** | Cande | 145040-37-5 | Sigma Aldrich | SML0245 | 2x10-5 | 4x10-6 | Yes<br>PPARγ ligand | > 98% | (Erbe et al. 2006) |
| **CL 316,243** | CL316 | 138908-40-4 | Sigma Aldrich | C5976 | 5x10-6 | 5x10-6 | No<br>β3 agonist | > 98% | --- |
| **Corticosterone** | Corti | 50-22-6 | Sigma Aldrich | 27840 | 2x10-6 | 2x10-6 | No<br>GR ligand | > 98% | --- |
| **Cyazofamid** | Cyazo | 120116-88-3 | Sigma Aldrich | 33874 | 4x10-5 | 2x10-5 | Potential | NA | (Auerbach et al. 2016) |
| **d-cis,trans-Allethrin** | Allet | 548-79-2 | Sigma Aldrich | 33396 | 2x10-5 | 1x10-5 | Potential | 97% | (Auerbach et al. 2016) |
| **Dexamethasone** | Dex-SP | 2392-39-4 | Sigma Aldrich | D1159 | 2x10-7 | 2x10-7 | No<br>GR ligand | > 98% | --- |
| **Di(2-ethylhexyl) phthalate** | DEHP | 117-81-7 | Sigma Aldrich | 36735 | 1x10-5 | 1x10-5 | No | > 99% | (Feige et al. 2007) |
| **Dibutyltin** | DBT | 683-18-1 | Sigma Aldrich | 205494 | 2x10-7 | 2x10-7 | Yes<br>PPARγ/RXR ligand | 96% | Hiromori et al., 2009 |
| **Diisononyl phthalate** | DINP | 28553-12-0 | Sigma Aldrich | 376663 | 1*10-5 | 1*10-5 | Yes | > 99% | (Zhang et al., 2019) |
| **Dioctyl sulfosuccinate sodium** | DOSS | 577-11-7 | Sigma Aldrich | 323586 | 5x10-6 | 5x10-6 | Potential | > 97% | (Temkin et al. 2016) |
| **Diphenyl phosphate** | DiPhPho | 838-85-7 | Sigma Aldrich | 850608 | 1x10-5 | 1x10-5 | Potential | 99% | (Cano-Sancho et al. 2017) |
| **Ethylene brassylate** | EtBra | 105-95-3 | Sigma Aldrich | W354309 | 1x10-5 | 1x10-5 | No evidence | > 95% | --- |
| **Fenthion** | Fenth | 55-38-9 | Sigma Aldrich | 36552 | 4x10-5 | 4x10-5 | Potential | > 99% | (Auerbach et al. 2016) |
| **Firemaster 550** | FM550 | Gift from Heather Stapleton, Duke | | | 10 ug/ml | 10 ug/ml | Yes<br>Induces PPARγ target genes and 3T3 L1 adipogenesis | NA | (Pillai et al. 2014) |
| **Fludioxonil** | Fludi | 131341-86-1 | Sigma Aldrich | 46102 | 2x10-5 | 2x10-6 | Potential | > 95% | (Auerbach et al. 2016) |
| **Honokiol** | Honok | 35354-74-6 | Sigma Aldrich | H4914 | 2x10-5 | 4x10-6 | Yes<br>PPARγ ligand | > 98% | (Atanasov et al. 2013) |
| **LG100268** | LG268 | 153559-76-3 | Sigma Aldrich | SML0279 | 1x10-7 | 1x10-7 | Yes<br>Activates PPARγ through RXR | > 98% | (Cesario et al. 2001) |

| | | | | | | | | | |
|---|---|---|---|---|---|---|---|---|---|
| **LG100754** | LG754 | 180713-37-5 | Tocris | 3831 | 2x10-7 | 2x10-7 | Yes<br>Activates PPARγ through RXR | > 99% | (Cesario et al. 2001) |
| **Magnolol** | Magno | 528-43-8 | Sigma Aldrich | M3445 | 2x10-5 | 2x10-5 | Yes<br>PPARγ ligand | > 95% | (Fakhurudin et al., 2010) |
| **MCC-555** | MCC555 | 161600-01-7 | Sigma Aldrich | SML0896 | 5x10-6 | 5x10-6 | Yes<br>PPARγ ligand | 98% | (Reginato et al. 1998) |
| **Melengestrol acetate** | Melen | 2919-66-6 | Sigma Aldrich | 73248 | 2x10-5 | 2x10-5 | No<br>PR ligand | > 97% | --- |
| **Mono-(2-ethyhexyl) tetrabromophthalate** | METBP | Synthesized by Asis Chemical | | | 1x10-5 | 1x10-5 | Yes<br>Activates PPARγ reporter and induces adipogenesis in multipotent stromal cells | 95% | (Watt and Schlezinger 2015) |
| **Mono(2-ethylhexyl) phthalate** | MEHP | 4376-20-9 | Sigma Aldrich | CDS01060 8 | 1x10-5 | 1x10-5 | Yes<br>PPARγ ligand | NA | (Feige et al. 2007) |
| **Monobenzyl phthalate** | MBzP | 2528-16-7 | Sigma Aldrich | 89505 | 1x10-5 | 1x10-5 | Yes<br>Activates PPARγ reporter | 95% | (Hurst and Waxman 2003) |
| **Mono-n-butyl phthalate** | MBuP | 131-70-4 | Sigma Aldrich | 30751 | 2x10-5 | 2x10-5 | Yes<br>Activates PPARγ reporter and induces 3T3 L1 adipogenesis | > 98% | (Hurst and Waxman 2003) |
| **n-Butylparaben** | BuPara | 94-26-8 | Sigma Aldrich | 54680 | 2x10-5 | 2x10-5 | Yes<br>Activates PPARγ reporter and induces 3T3 L1 adipogenesis | > 99% | (Hu et al. 2013) |
| **N-nitro-2-imidazolidinimine** | Imida | 138261-41-3 | Sigma Aldrich | 37894 | 1x10-5 | 1x10-5 | No evidence | > 99% | --- |
| **nTZDpa** | nTZDpa | 118414-59-8 | Sigma Aldrich | SML0616 | 1x10-6 | 1x10-6 | Yes<br>PPARγ ligand | > 98% | (Berger et al. 2003) |
| **Perfluorooctanesulfonic acid** | PFOS | 2795-39-3 | Sigma Aldrich | 77282 | 4x10-5 | 4x10-5 | Potential | NA | (Takacs and Abbott 2007) |
| **Perfluorooctanoic acid** | PFOA | 335-67-1 | Sigma Aldrich | 33824 | 1x10-5 | 1x10-5 | Potential | > 99% | (Takacs and Abbott 2007) |
| **Pioglitazone hydrochloride** | Piogl | 112529-15-4 | Sigma Aldrich | E6910 | 1x10-5 | 1x10-5 | Yes<br>PPARγ ligand | > 98% | (Gimble et al. 1996) |
| **Prallethrin** | Prall | 23031-36-9 | Sigma Aldrich | 32917 | 1x10-5 | 1x10-5 | Potential | > 95% | (Auerbach et al. 2016) |
| **Pregnenolone 16α-carbonitrile** | Pregn | 1434-54-4 | Sigma Aldrich | P0543 | 1x10-5 | 1x10-5 | No<br>PXR ligand | > 97% | --- |

| | | | | | | | | | |
|---|---|---|---|---|---|---|---|---|---|
| **Propylparaben** | ProPara | 94-13-3 | Sigma Aldrich | P53357 | 1x10-5 | 1x10-5 | Yes Activates PPARγ reporter and induces 3T3 L1 adipogenesis | > 99% | (Pereira-Fernandes et al. 2013) |
| **Protectin D1** | Prote | 660430-03-5 | Cayman Chemical | 10008128 | 2x10-6 | 2x10-6 | Yes PPARγ ligand | > 98% | (Muralikumar et al. 2017) |
| **Quinoxyfen** | Quino | 124495-18-7 | Sigma Aldrich | 46439 | 1x10-5 | 1x10-5 | Potential | NA | (Auerbach et al. 2016) |
| **Resolvin-E1** | Resol | 552830-51-0 | Cayman Chemical | 10007848 | 2x10-6 | 2x10-6 | Yes PPARγ ligand | > 95% | (Muralikumar et al. 2017) |
| **Roscovitine** | Rosco | 186692-46-6 | Sigma Aldrich | R7772 | 4x10-5 | 4x10-6 | Yes Inhibits CDK5, preventing PPARγ phosphorylation | > 98% | (Wang et al. 2016) |
| **Rosiglitazone** | Rosig | 122320-73-4 | Cayman Chemical | 71740 | 1x10-6 | 1x10-6 | Yes PPARγ ligand | > 98% | (Lehmann et al. 1995) |
| **S26948** | S26948 | 353280-43-0 | Sigma Aldrich | SML0510 | 2x10-6 | 2x10-6 | Yes PPARγ ligand | > 98% | (Carmona et al. 2007) |
| **Sodium arsenite** | Arsen | 7784-46-5 | Sigma Aldrich | S7400 | 4x10-7 | 4x10-7 | No, Reduces adipogenesis | > 90% | (Wauson et al. 2002) |
| **Sodium tungstate** | Tungs | 10213-10-2 | Sigma Aldrich | 14304 | 2x10-5 | 2x10-5 | No, Reduces adipogenesis | > 99% | (Carmona et al. 2009) |
| **SR1664** | SR1664 | 1338259-05-4 | Sigma Aldrich | SML0636 | 1x10-6 | 1x10-6 | Yes PPARγ ligand | 98% | (Choi et al. 2011) |
| **T0901317** | T1317 | 293754-55-9 | Sigma Aldrich | T2320 | 1x10-6 | 1x10-6 | No, LXR ligand | > 98% | --- |
| **T0070907** | T007 | 313516-66-4 | Sigma Aldrich | T8703 | 4x10-5 | 8x10-6 | Yes PPARγ antagonist | > 98% | (Lee et al. 2002) |
| **Tebuconazole** | Tebuc | 107534-96-3 | Sigma Aldrich | 32013 | 2x10-5 | 2x10-5 | Potential | NA | (Auerbach et al. 2016) |
| **Telmisartan** | Telmi | 144701-48-4 | Sigma Aldrich | T8949 | 2x10-5 | 2x10-5 | Yes PPARγ ligand | > 98% | (Yamagishi and Takeuchi 2005) |
| **Tesaglitazar** | Tesag | 251565-85-2 | Sigma Aldrich | SML1369 | 5x10-6 | 5x10-6 | Yes PPARγ ligand | > 98% | (Ljung et al. 2002) |
| **Tolylfluanid** | Tolyl | 731-27-1 | Sigma Aldrich | 32060 | 2x10-7 | 2x10-7 | No GR ligand | > 99% | (Regnier et al. 2015) |
| **Tonalide** | Tonal | 21145-77-7 | Sigma Aldrich | W526401 | 4x10-6 | 4x10-6 | Potential | > 98% | (Pereira-Fernandes et al. 2013) |
| **Tributyl phosphate** | TBuP | 126-73-8 | Sigma Aldrich | 240494 | 2x10-5 | 2x10-5 | Yes PPARγ ligand | > 99% | (Fang et al. 2015a) |

| | | | | | | | | | |
|---|---|---|---|---|---|---|---|---|---|
| **Tributyltin** | TBT | 1461-22-9 | Sigma Aldrich | T50202 | 8x10-8 | 8x10-8 | Yes PPARγ/RXR ligand | 96% | (Grun et al. 2006) |
| **Triflumizole** | Trifl | 68694-11-1 | Sigma Aldrich | 32611 | 2x10-5 | 2x10-5 | Yes Activates PPARγ reporter and induces 3T3 L1 adipogenesis | NA | (Li et al. 2012) |
| **Triphenyl phosphate** | TPhP | 115-86-6 | Sigma Aldrich | 241288 | 2x10-5 | 1x10-5 | Yes PPARγ ligand | > 99% | (Pillai et al. 2014) |
| **Triphenyl phosphite** | TPhPhi | 101-02-0 | Sigma Aldrich | T84654 | 1x10-5 | 1x10-5 | Potential | 97% | (Fang et al. 2015a) |
| **Triphenylphosphine oxide** | TPhPho Ox | 791-28-6 | Sigma Aldrich | T84603 | 1x10-5 | 1x10-5 | Potential | 98% | (Hiromori et al. 2016) |
| **Triphenyltin** | TPhT | 639-58-7 | Sigma Aldrich | 245712 | 8x10-8 | 8x10-8 | Yes PPARγ/RXR ligand | 98% | (Hiromori et al., 2009) |
| **Tris(1,3-dichloro-2-propyl) phosphate** | TDCPP | 13674-87-8 | Sigma Aldrich | 32951 | 2x10-5 | 2x10-5 | Potential | NA | (Fang et al. 2015b) |
| **Tris(1-chloro-2-propyl) phosphate** | TCCP | 13674-87-5 | Sigma Aldrich | 32952 | 1x10-5 | 1x10-5 | Potential | NA | (Fang et al. 2015b) |
| **Troglitazone** | Trogl | 97322-87-7 | Sigma Aldrich | T2573 | 5x10-6 | 5x10-6 | Yes PPARγ ligand | > 98% | (Lambe and Tugwood 1996) |

*NA = Not Available

[a] Toxicity was assessed by microscopic inspection. The concentration was reduced by two fold until a non-toxic, maximal concentration was identified. Maximal concentrations were determined in an N=4.

[b] "Yes" indicates there is experimental evidence of modification of PPARγ activity, including PPARγ binding assays, coactivator recruitment or computational modeling (definitive ligands), PPARγ-driven reporter assays (at least 25% of the rosiglitazone-induced maximum), expression of PPARγ target genes and/or differentiation of 3T3 L1 or multipotent stromal cells into adipocytes in the absence of a known PPARγ ligand. Chemicals that changed the expression of PPARγ (e.g., PCB126 and DDE) were not considered to be ligands or modifiers. "No" indicates that the chemical was chosen based on the fact that it is known to be a specific ligand of another receptor. "No evidence" indicates that this chemical has not been tested for PPARγ activation but is structurally dissimilar to known classes of PPARγ ligands. "Potential" indicates that the chemical was identified in other screening approaches or by the ToxPi designed to identify chemicals in the ToxCast dataset that have potential to be PPARγ ligands/modifiers.

[c] Abbreviations: AhR, aryl hydrocarbon receptor; ER, estrogen receptor; GR, glucocorticoid receptor; PR, progesterone receptor; PXR, pregnane X receptor; RAR, retinoic acid receptor; RXR, retinoid X receptor

**Table S2. Mouse (M) and human (H) primer sequences for reverse transcription qPCR.**

| GENE SYMBOL | FORWARD | REVERSE | ANNEALING TEMP. ° C |
|---|---|---|---|
| *M-Rn18s* | GTAACCCGTTGAACCCCATT | CCATCCAATCGGTAGTAGCG | 55 |
| *M-B2m* | CTGCTACGTAACACAGTTCCACCC | CATGATGCTTGATCACATGTCTCG | 55 |
| *M-Cidec* | AGGCCCTGTCGTGTTAGCAC | CATGATGCCTTTGCGAACCT | 55 |
| *M-Cidea* | TGCTCTTCTGTATCGCCCAGT | GCCGTGTTAAGGAATCTGCTG | 55 |
| *M-Elovl3* | TCCGCGTTCTCATGTAGGTCT | GGACCTGATGCAACCCTATGA | 55 |
| *M-Fabp4* | AGCCCAACATGATCATCAGC | TTTCCATCCCACTTCTGCAC | 55 |
| *M-Plin1* | GGGACCTGTGAGTGCTTCC | GTATTGAAGAGCCGGGATCTTTT | 55 |
| *M-Pgc1a* | AACAAGCACTTCGGTCATCCCTG | TTACTGAAGTCGCCATCCCTTAG | 55 |
| *M-Pparg2* | TGGGTGAAACTCTGGGAGATTC | AATTTCTTGTGAAGTGCTCATAGGC | 55 |
| *M-Rip140* | AGAACGCACATCAGGTGGCA | GATGGCCAGACACCCCTTTG | 55 |
| *M-Adipoq* | GCACTGGCAAGTTCTACTGCAA | GTAGGTGAAGAGAACGGCCTTGT | 55 |
| *M-Ucp1* | ACTGCCACACCTCCAGTCATT | CTTTGCCTCACTCAGGATTGG | 55 |
| *M-Acaa2* | TAACGAGGCTGGCTACTTCAA | AGGGGCGTGAAGTTATGTTTT | 55 |
| *H-RPL27* | GTGAAAGTGTATAACTACAATCACC | TCAAACTTGACCTTGGCCT | 58 |
| *H-B2M* | GCTATCCAGCGTACTCCAAAG | CACACGGCAGGCATACTC | 58 |
| *H-CIDEC* | GGGATACAGTGTTCATGGTCCT | TCAATCTTCTTGGCAGGCTTATG | 55 |
| *H-CIDEA* | GGCAGGTTCACGTGTGGATA | GAAACACAGTGTTTGGCTCAAGA | 60 |

**Table S3. Metabolic parameters included and excluded in human transcriptome analysis.**

| PARAMETERS INCLUDED | PARAMETERS EXCLUDED |
|---|---|
| Fat free mass % | Body mass index (kg/m2) |
| Fasting Plasma parameters | Waist-to-hip ratio |
| Free fatty acid (mmol/l) | Waist circumference (cm) |
| Total triglycerides (mmol/l) | Hip circumference (cm) |
| LDL cholesterol (mmol/l) | Plasma total fatty acids (mmol/l) |
| HDL cholesterol (mmol/l) | Plasma total cholesterol (mmol/l) |
| Adiponectin (ug/ml) | Matsuda composite insulin sensitivity index |
| Glucose (mmol/l) | HOMA-IR |
| Insulin (mU/l) | Systolic blood pressure (mm Hg) |
| Proinsulin (pmol/l) | Diastolic blood pressure (mm Hg) |
| Glycated HbA1c (%) | Glomerular filtration rate |
| High sensitivity C-reactive protein (mg/l) | |
| Interleukin-1 receptor antagonist (pg/ml) | |

**Excel File 1. Detailed results from random forest classification.**

**Excel File 2. Detailed annotation of clustering results for individual modules.**

**Excel File 3. Supporting numerical data from all cell culture experiments.**

**Excel File 4. Detailed results of partial correlation analysis of clinical measurements and projections of chemical taxonomy gene signatures onto human adipose gene expression.**

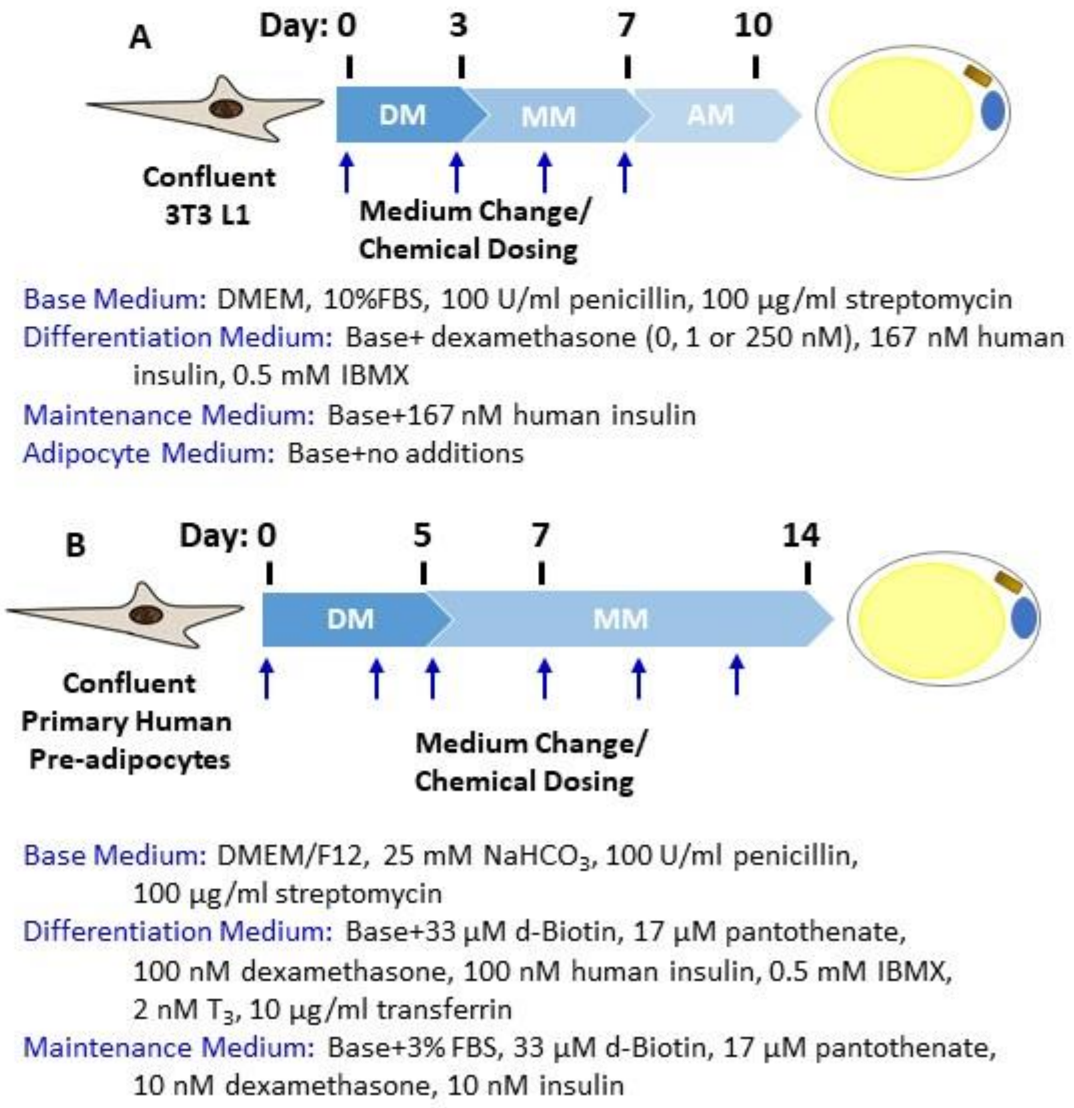


**Figure S1. Differentiation and dosing protocols for 3T3-L1 cells (A) and primary human preadipocytes (B).**

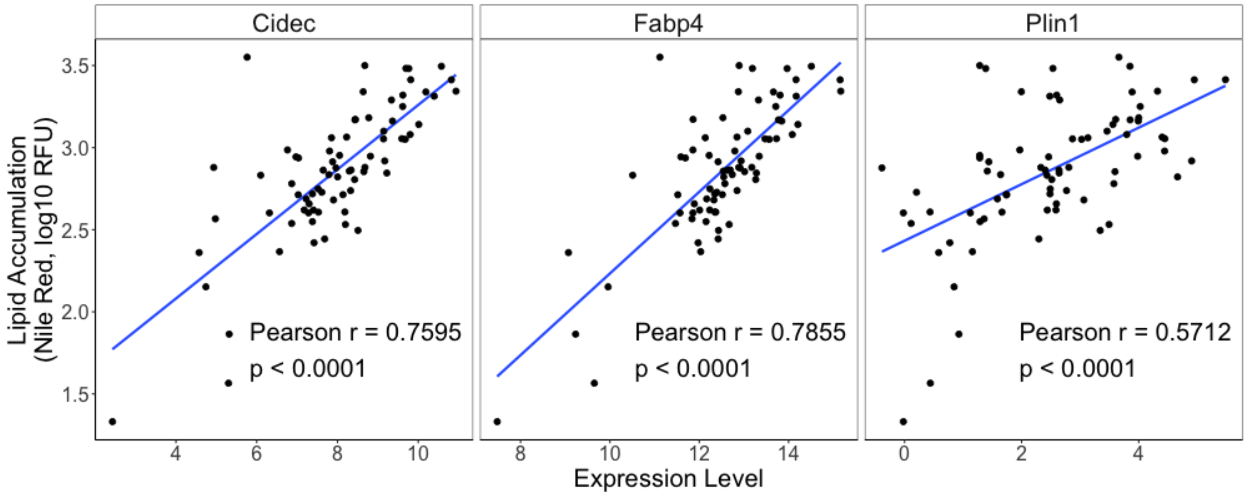


**Figure S2. Correlation of lipid accumulation with *Cidec, Fabp4, and Plin1* expression in differentiated and treated 3T3-L1 pre-adipocytes.** Confluent 3T3 L1 cells were differentiated using a standard hormone cocktail for 10 days. During differentiation, cells were treated with vehicle (Vh, 0.1% DMSO, final concentration) or test chemical (**Table S1**). On days 3, 5, and 7 of differentiation, the medium was replaced and the cultures re-dosed. Following 10 days of differentiation and dosing, cells were analyzed for lipid accumulation by Nile Red staining (Data are from Figure 1) and *gene* expression by 3'DGE. Each point represents the mean data for each chemical, (n=2-4). The least squares linear model estimate is shown in blue.

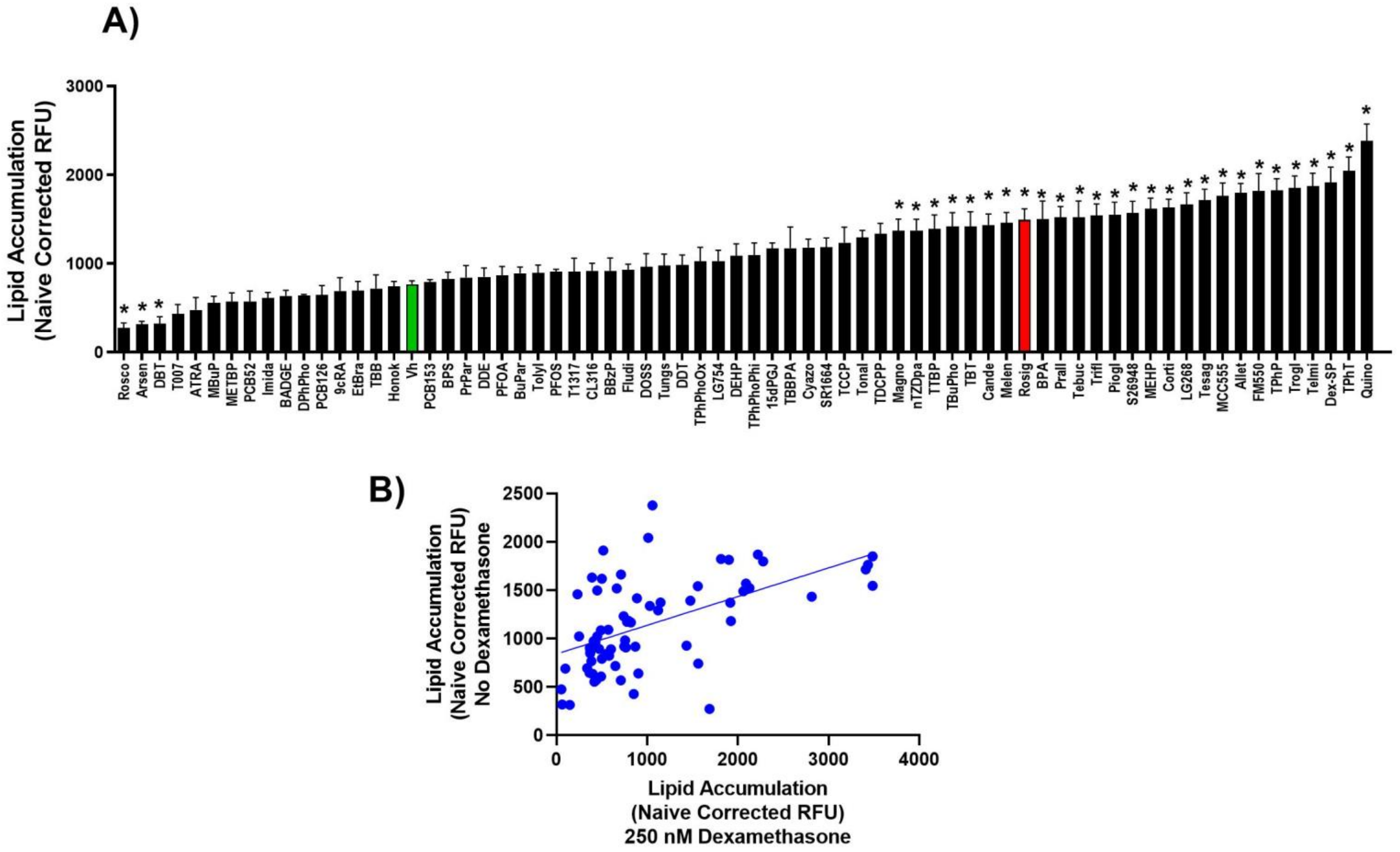


**Figure S3. Lipid accumulation in differentiated and treated 3T3-L1 pre-adipocytes in the absence of dexamethasone.**

Confluent 3T3 L1 cells were differentiated using a standard hormone cocktail for 10 days, with the exception of using no dexamethasone. During differentiation, cells were treated with vehicle (Vh, 0.2% DMSO, final concentration), rosiglitazone (positive control, 100 nM) or test chemical (**Table S1**). On days 3, 5, and 7 of differentiation, the medium was replaced and the cultures re-dosed. Following 10 days of differentiation and dosing, cells were analyzed for lipid accumulation by Nile Red staining. **A)** Nile Red staining induced by individual chemicals. Nile Red fluorescence was normalized by subtracting the fluorescence measured in naïve pre-adipocyte cultures within each experiment and reported as "Naïve Corrected RFU." Data are presented as mean ± SE (n=4). Statistically different from Vh-treated (highlighted in green) (*$p<0.05$, ANOVA, Dunnett's). **B)** Correlation between lipid accumulation induced in 3T3 L1 cells differentiated in the presence (data are from Figure 1) and absence of dexamethasone. Numerical data are provided in **Excel File 3**. Pearson's r = 0.5429 ($p<0.0001$).

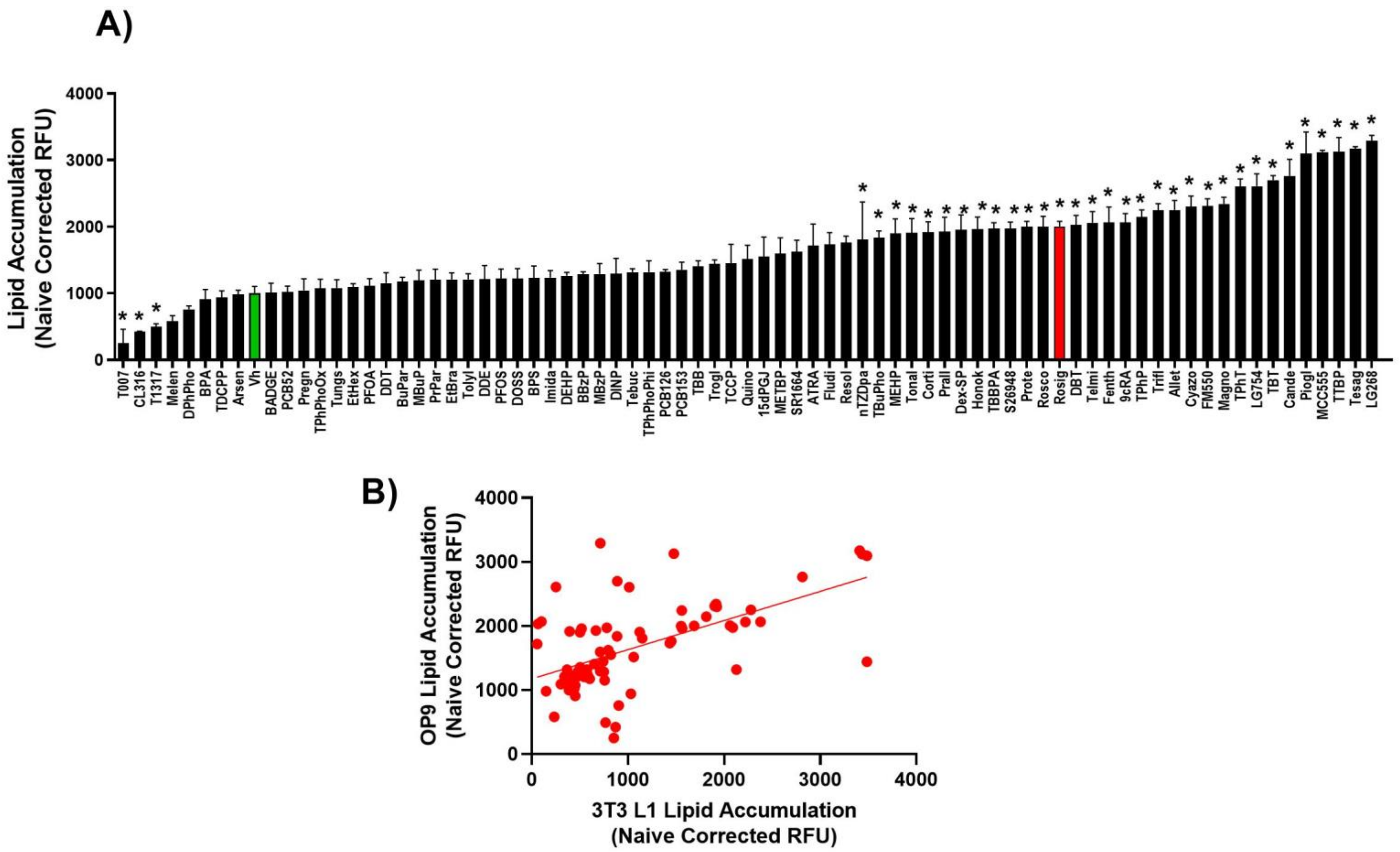


**Figure S4. Lipid accumulation in differentiated and treated OP9 pre-adipocytes.**

Confluent OP9 cells were differentiated using a standard hormone cocktail for 10 days, with the exception of using 125 nM dexamethasone. During differentiation, cells were treated with vehicle (Vh, 0.2% DMSO, final concentration), rosiglitazone (positive control, 100 nM) or test chemical (**Table S1**). On days 3, 5, and 7 of differentiation, the medium was replaced and the cultures re-dosed. Following 10 days of differentiation and dosing, cells were analyzed for lipid accumulation by Nile Red staining. **A)** Nile Red staining induced by individual chemicals. Nile Red fluorescence was normalized by subtracting the fluorescence measured in naïve pre-adipocyte cultures within each experiment and reported as “Naïve Corrected RFU.” Data are presented as mean ± SE (n=4). Statistically different from Vh-treated (highlighted in green) (*p<0.05, ANOVA, Dunnett’s). **B)** Correlation between lipid accumulation induced in 3T3 L1 cells differentiated in the presence of dexamethasone (data are from Figure 1) and in OP9 cells. Numerical data are provided in **Excel File 3**. Pearson’s r = 0.5768 (p<0.0001).

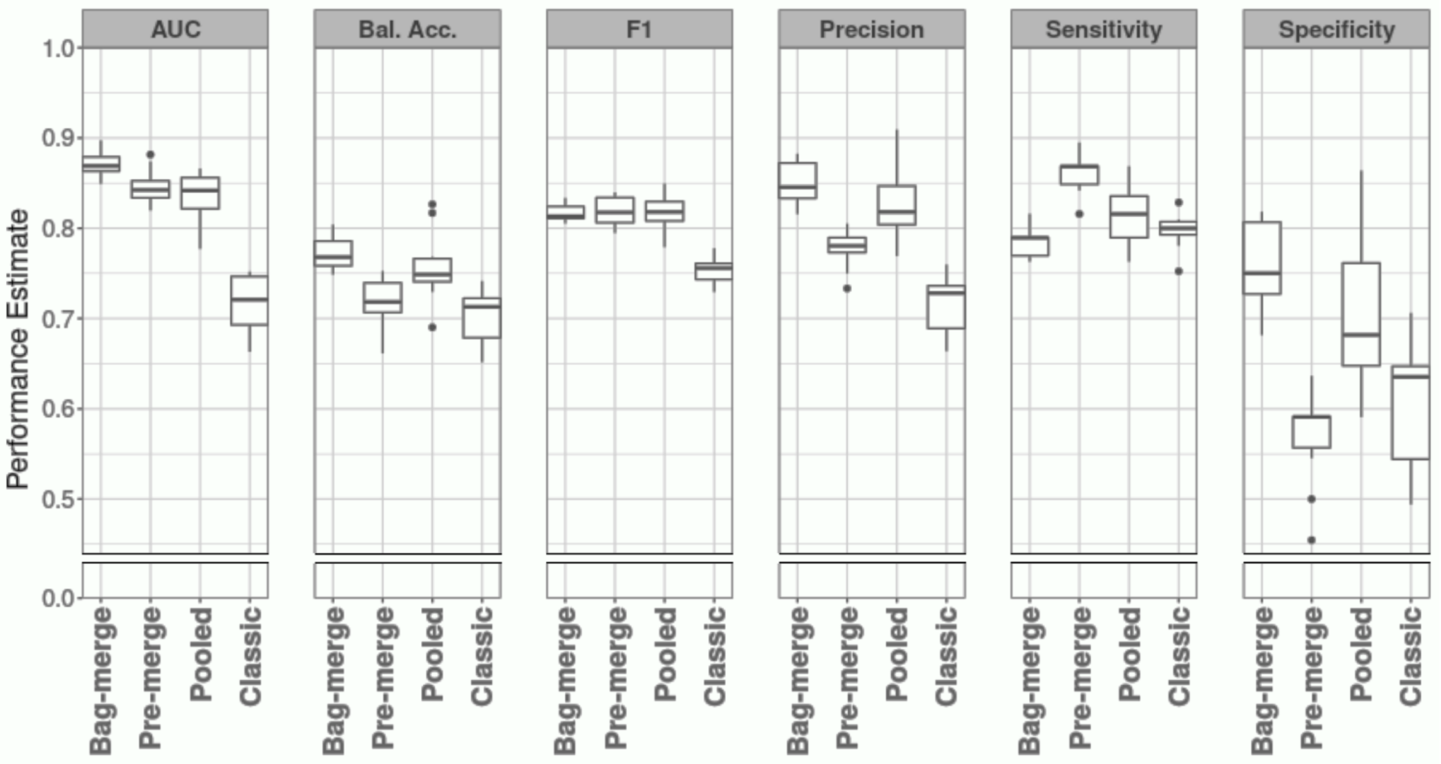


**Figure S5. Performance comparison of random forest methods.**
Boxplots of performance estimates for repeated 10-fold cross validation for each of the four random forest methods considered for classifying PPARγ activity modifying compounds from high-throughput gene expression profiles of chemically treated 3T3 L1 cells. Abbreviated metrics shown include: area-under the curve (AUC), balanced accuracy (Bal. Acc.), F1-score (F1). Besides AUC, for all performance metrics besides, an appropriate classification threshold for predicting PPARγ activity modifying compound labels in each test set was estimated based on out-of-bag voting of their corresponding training set. Sensitivity and specificity are the proportions of identified known PPARγ activity modifying compounds and known non-PPARγ activity modifying compounds, respectively, out of the total number of each label in the data. Bal. Acc. is the mean of sensitivity and specificity. Precision is the proportion of accurately identified known PPARγ activity modifying compounds out of all predicted PPARγ activity modifying compounds. F1 is the harmonic mean of sensitivity and precision. Finally, AUC is the integral of sensitivity and specificity across every possible classification threshold. The distribution of each method/metric combination is based on 10 repetitions of cross validation (i.e. N=10). The full set of performance estimates for each repetition, performance metric, and classification procedure considered are shown in **Excel File 3**. The midline, box limits, and whiskers show the median, upper/lower quartile, and minimum/maximum of each distribution cutoff at a distance of 1.5 * the interquartile range from their closest box limit, respectively. Individual points indicate values which extend beyond the whisker limits. The horizontal gap in each plot indicates that, while the range of possible estimates for each performance metric is between 0.0 and 1.0, all estimates fell between 0.45 and 1.0.

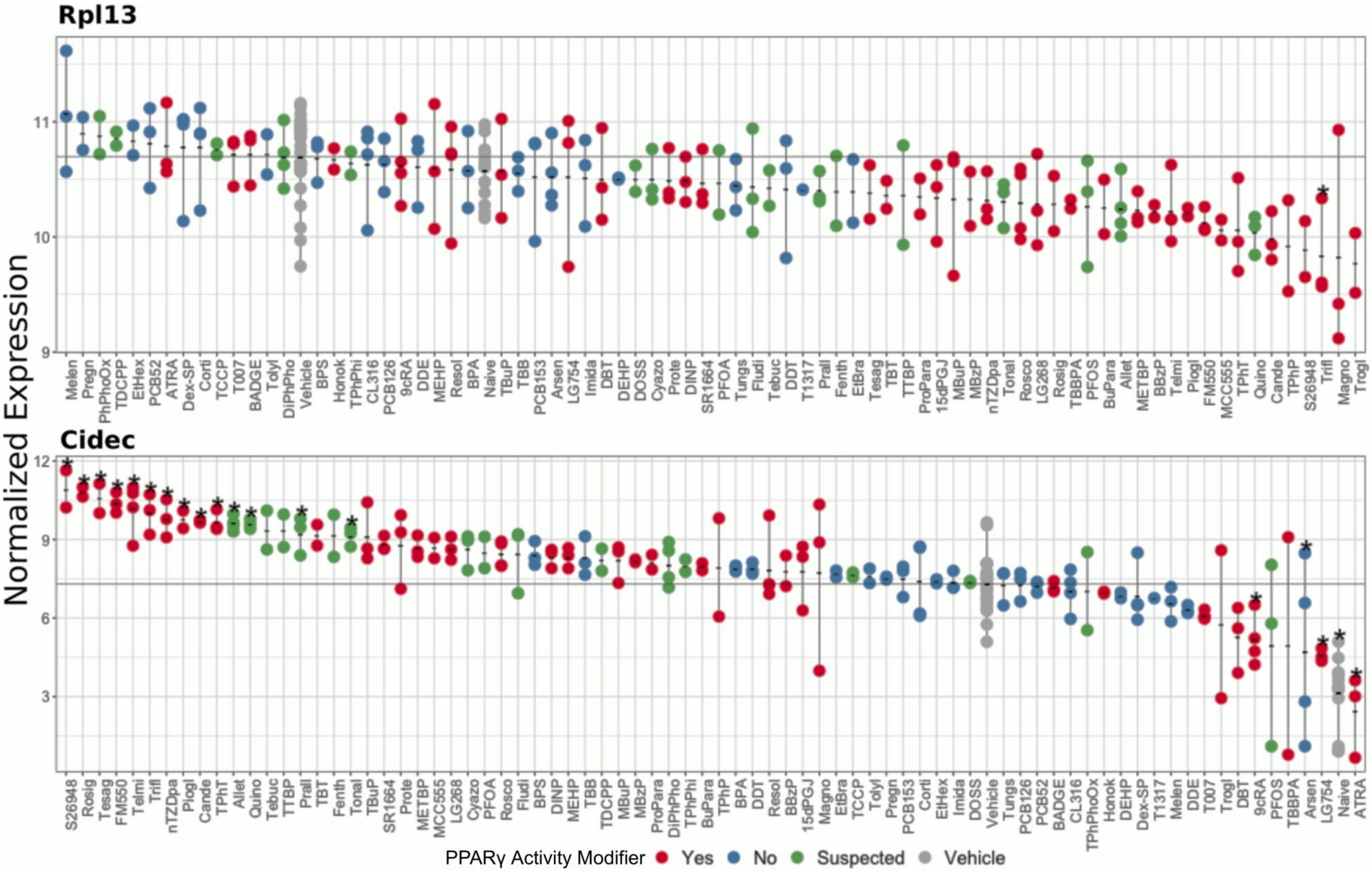
Rpl13
Cidec
Normalized Expression
11
10
9
12
9
6
3
Melen Pregn PhPhoOx TDCPP EtHex PCB52 ATRA Dex-SP Corti TCCP T007 BADGE Tolyl DiPhPho Vehicle BPS Honok TPhPhi CL316 PCB126 9cRA DDE MEHP Resol BPA Naive TBuP TBB PCB153 Arsen LG754 Imida DBT DEHP DOSS Cyazo Prote DINP SR1664 PFOA Tungs Fludi Tebuc DDT T1317 Prall Fenth EtBra Tesag TBT TTBP ProPara 15dPGJ MBuP MBzP nTZDpa Tonal Rosco LG268 Rosig TBBPA PFOS BuPara Allet METBP BBzP Telmi Piogl FM550 MCC555 TPhT Quino Cande TPhP S26948 Trifl Magno Trogl
S26948 Rosig Tesag FM550 Telmi Trifl nTZDpa Piogl Cande TPhT Allet Quino Tebuc TTBP Prall TBT Fenth Tonal TBuP SR1664 Prote METBP MCC555 LG268 Cyazo PFOA Rosco Fludi BPS DINP MEHP TBB TDCPP MBuP MBzP ProPara DiPhPho TPhPhi BuPara TPhP BPA DDT Resol BBzP 15dPGJ Magno EtBra TCCP Tolyl Pregn PCB153 Corti EtHex Imida DOSS Vehicle Tungs PCB126 PCB52 BADGE CL316 TPhPhoOx Honok DEHP Dex-SP T1317 Melen DDE T007 Trogl DBT 9cRA PFOS TBBPA Arsen LG754 Naive ATRA
PPARγ Activity Modifier
Yes
No
Suspected
Vehicle

**Figure S6. Classification Results (Distributions of individual genes).** Confluent 3T3 L1 cells were differentiated using a standard hormone cocktail for 10 days. During differentiation, cells were treated with 0.1% DMSO, final concentration (*vehicle*), test chemical, or were untreated (*naive*). On days 3, 5, and 7 of differentiation, the medium was replaced and the cultures re-dosed. Following 10 days of differentiation and dosing, cells were analyzed for *gene* expression by 3'DGE. The labels, "Yes", "No", and "Potential", indicate test chemicals predetermined to be known PPARγ activity modifying compounds, known <u>non</u>-PPARγ activity modifying compounds, and Potential PPARγ activity modifying compounds, respectively, based on previous studies (See **Table S1**). *Rpl13* and *Cidec* demonstrated the greatest predictive value for classifying PPARγ activity modifying compounds based on their Gini importance estimates from random forest modeling (Breiman 2001). Each point indicates sample-specific expression values, normalized by batch correction and Trimmed Mean of M-values (TMM) transformation, performed in R (v 3.4.3) using *ComBat* (v 3.26.0) (Leek et al. 2012) and *limma* (v 3.34.9) (Ritchie et al. 2015), respectively. The range and mean expression of the biological replicates of each treatment is indicated by a vertical and short horizontal line, respectively. Mean expression across all *vehicle* samples is shown as a horizontal line spanning the plot. Exposures which have statistically significant different means from *vehicle* (FDR Q-value < 0.05) are highlighted with an asterisk. For test chemicals sample sizes range from N=2-4. For *vehicle* and *naive* sample sizes are N=25 and N=15, respectively.

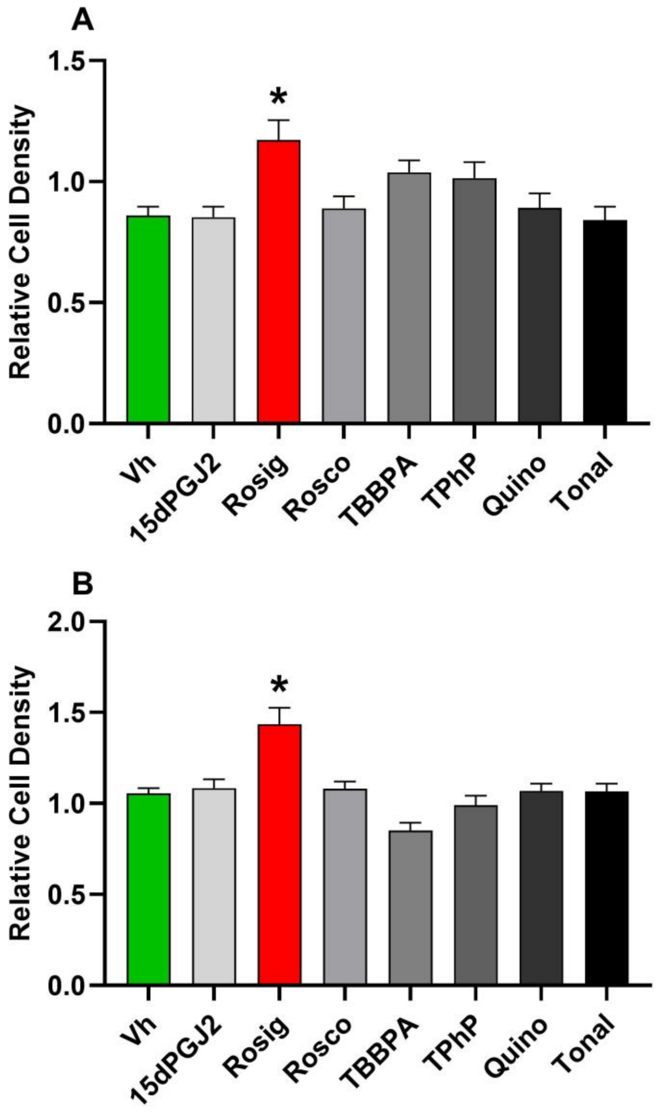


**Figure S7. Cell number analyses in the differentiated and treated 3T3-L1s.** Confluent 3T3 L1 cells were differentiated using a standard hormone cocktail with 1 nM dexamethasone **(A)** and 250 nM dexamethasone **(B)** for 10 days. During differentiation, cells were treated with Vh (0.1% DMSO, final concentration), rosiglitazone (Rosig, 200 nM (A), 1 μM (B)), roscovitine (Rosco, 4 μM), 15dPGJ2 (500 nM (A), 1 μM (B)), TBBPA (20 μM) and TPhP (10 μM). On days 3, 5, and 7 of differentiation, the adipocyte maintenance medium was replaced and the cultures re-dosed. Cells were incubated for a total of 10 days of differentiation. To assess cell number, cells were stained with Janus green stain. Absorbance in experimental wells was normalized by dividing by the absorbance measured in naïve pre-adipocyte cultures within the experiment and reported as "Relative Cell Density." Numerical data are provided in **Excel File 3**. Data are presented as means ± SE (n=8). Statistically different from Vh-treated (**p<0.01, ANOVA, Dunnett's).

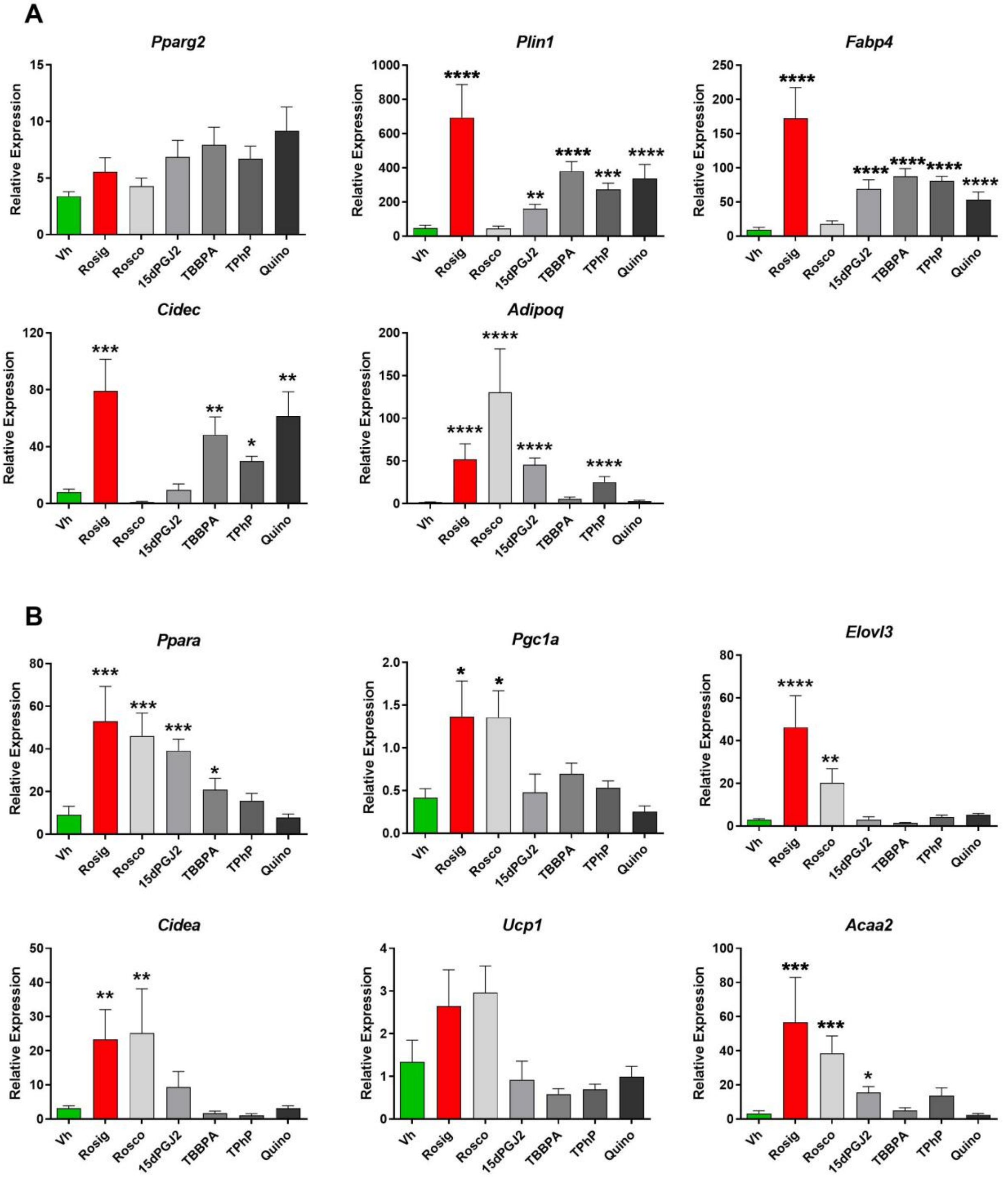


**Figure S8.** White and brite gene expression in differentiated and treated 3T3-L1 adipocytes. Confluent 3T3 L1 cells were differentiated using a standard hormone cocktail with 250 nM dexamethasone for 10 days. During differentiation, cells were treated with vehicle (Vh, 0.2% DMSO, final concentration), rosiglitazone (Rosig, 200 nM), roscovitine (Rosco, 4 μM), 15dPGJ2 (500 nM), TBBPA (20 μM), TPhP (10 μM) and quinoxyfen (10 μM). On days 3, 5, and 7 of differentiation, the adipocyte maintenance medium was replaced and the cultures re-dosed. Following 10 days of differentiation and dosing, cells were analyzed for gene expression by RT-qPCR. **(A)** Genes related to white adipogenesis. **(B)** Genes related to brite adipogenesis. Gene expression levels were normalized to the geometric mean of the expression levels of *B2m* and *Rn18s* and expressed as "Relative Expression" in comparison to naïve, pre-adipocyte cultures using the Pfaffl method. Numerical data are provided in **Excel File 3**. Data are presented as mean ± SE of n=6 independent experiments. Statistically different from Vh-treated (highlighted in green) (*$p<0.05$, **$p<0.01$, ***$p<0.001$, ****$p<0.0001$, ANOVA, Dunnett’s).

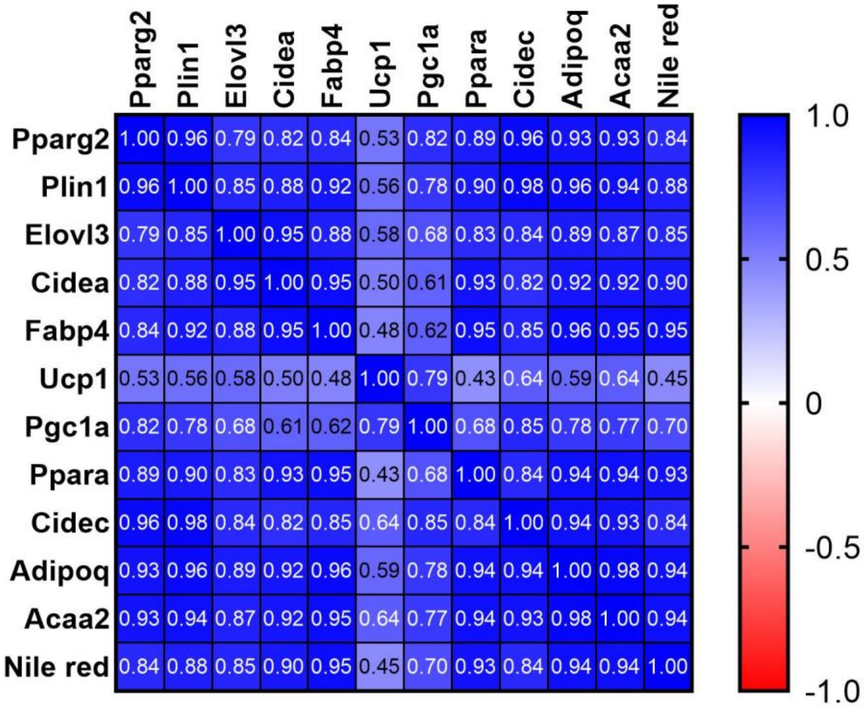


**Figure S9.** Spearman correlation analysis of lipid accumulation (Nile Red) and gene expression. Confluent 3T3 L1 cells were differentiated using a standard hormone cocktail with 1 nM dexamethasone for 10 days and analyzed for adipocyte differentiation by staining for lipids with Nile Red (shown in Figures 6 and 11) and gene expression (shown in Figures 7 and 11).

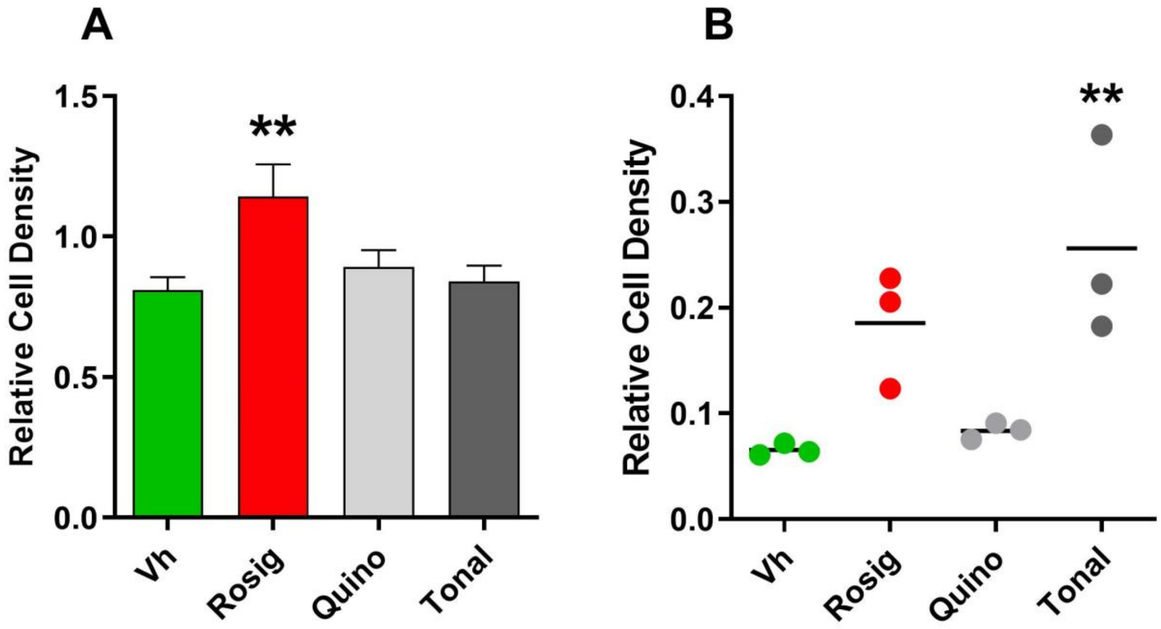


**Figure S10.** Cell number analyses in the differentiated and quinoxyfen and tonalide treated 3T3-L1s and human primary preadipocytes. **(A)** Confluent 3T3 L1 cells were differentiated using a standard human adipocyte hormone cocktail for 10 days. During differentiation, cells were treated with Vh (0.1% DMSO, final concentration), rosiglitazone (Rosig, 200 nM), quinoxyfen (Quino, 10 µM) or tonalide (Tonal, 4 µM). On days 3, 5, and 7 of differentiation, the adipocyte maintenance medium was replaced and the cultures re-dosed. Cells were incubated for a total of 10 days of differentiation. Data are presented as means ± SE (n=8). **(B)** Confluent primary human preadipocytes were differentiated using a standard hormone cocktail for 14 days. During differentiation, cells were treated with Vh (0.1% DMSO, final concentration), rosiglitazone (Rosig, 4 µM), quinoxyfen (Quino, 4 µM) or tonalide (4 µM). On days 3, 5, 7, 10, and 12 of differentiation, the medium was replaced and the cultures re-dosed. Following 14 days of differentiation and dosing, cultures were analyzed for relative cell density using JANUS Green staining. Absorbance in experimental wells was normalized by dividing by the absorbance measured in naïve pre-adipocyte cultures within the experiment and reported as "Relative Cell Density." Data are presented as mean ± SE (n=3, each n is from adipocytes from an individual). Statistically different from Vh-treated (**$p<0.01$, ANOVA, Dunnett's).
